# MLL3/MLL4 methyltransferase activities control early embryonic development and embryonic stem cell differentiation in a lineage-selective manner

**DOI:** 10.1101/2020.09.14.296558

**Authors:** Guojia Xie, Ji-Eun Lee, Anna D. Senft, Young-Kwon Park, Shreeta Chakraborty, Joyce J. Thompson, Chengyu Liu, Todd S. Macfarlan, Pedro P. Rocha, Weiqun Peng, Kai Ge

## Abstract

H3K4me1 methyltransferases MLL3 (KMT2C) and MLL4 (KMT2D) are critical for enhancer activation, cell differentiation and development. However, roles of MLL3/4 enzymatic activities and MLL3/4-mediated enhancer H3K4me1 in these processes remain unclear. Here, we report that constitutive elimination of both MLL3 and MLL4 enzymatic activities leads to gastrulation failure and early embryonic lethality in mice. However, selective elimination of MLL3/4 enzymatic activities in embryonic, but not extraembryonic, lineages leaves gastrulation largely intact. Consistently, embryonic stem cells (ESCs) lacking MLL3/4 enzymatic activities can differentiate towards the three embryonic germ layers but show aberrant differentiation to extraembryonic endoderm and trophectoderm. The failure in extraembryonic endoderm differentiation can be attributed to markedly reduced enhancer-binding of the lineage-determining transcription factor GATA6. Furthermore, we show that MLL3/4-catalyzed H3K4me1 is largely dispensable for enhancer activation during ESC differentiation. Together, our findings suggest a lineage-selective, but enhancer activation-independent, role of MLL3/4 methyltransferase activities in early embryonic development and embryonic stem cell differentiation.

## Introduction

Enhancers are *cis*-regulatory DNA elements recognized by transcription factors (TFs). They communicate with promoters to regulate cell-type-specific gene expression and cell identity. In-depth epigenomic research has uncovered chromatin signatures of enhancers. Primed enhancers, which are marked by H3K4me1, become activated with the addition of H3K27ac^1^. The activation of enhancers during cell fate transition is enabled by several epigenomic regulators sequentially recruited by lineage determining TFs, which include H3K4me1 methyltransferases MLL3 (KMT2C) and MLL4 (KMT2D) followed by H3K27ac acetyltransferases CBP and p300^2–5^. However, the exact role of these epigenomic regulators themselves versus the histone modifications they catalyze, particularly MLL3/4 versus MLL3/4-catalyzed H3K4me1, in enhancer activation has remained elusive.

MLL3 and MLL4 are members of the Set1-like family of mammalian H3K4 methyltransferases and are responsible for catalyzing H3K4me1^3, 6^. MLL3 and MLL4 are the largest known nuclear proteins with 4,903 and 5,588 amino acids in mice, respectively. They associate with the WRAD (WDR5, RBBP5, ASH2L, DPY30) subcomplex as well as NCOA6, UTX, PA1, and PTIP in a large multi-subunit complex^7, 8^. Enzymatic activities of MLL3/4 are conferred by the C-terminal SET domain, which is also required for maintaining their protein stability^9, 10^. Consistent with their critical roles in enhancer activation, MLL3/4 are broadly required for animal development and cell differentiation^6^. In humans, *MLL3/4* are frequently mutated in developmental diseases and cancers^6^. However, the functional roles of MLL3/4 enzymatic activities in animal development and cell differentiation are poorly understood.

The early development of mouse embryos from fertilization to gastrulation involves an orchestrated series of lineage specification events^11^. At the time of implantation, three cell lineages have been specified in the blastocyst: epiblast, primitive endoderm and trophectoderm. During gastrulation, the epiblast gives rise to the three embryonic germ layers: ectoderm, mesoderm and definitive endoderm. They contain progenitors of all fetal tissues. Primitive endoderm develops into visceral endoderm and parietal endoderm, which collectively are known as extraembryonic endoderm (ExEn). Together with trophectoderm and its derivatives, these extraembryonic tissues are critical for early embryonic development in mammals^12, 13^. Embryonic stem cells (ESCs) are self-renewing pluripotent cells that are derived from the inner cell mass of preimplantation-stage blastocysts^14^. ESCs can differentiate and organize into three-dimensional cavitated structures called embryoid bodies (EBs). EB differentiation of ESCs is a valuable *in vitro* model recapitulating major processes of early embryonic development, during which all three germ layers form^15^. Under the treatment of defined factors, ESCs can also differentiate homogeneously into specific cell types such as neurons and ExEn cells^16, 17^.

We previously reported that MLL3/4 proteins are required for enhancer activation and ESC differentiation but largely dispensable for ESC identity maintenance^4^. A recent study showed that MLL3/4-catalyzed H3K4me1 is mostly dispensable for maintaining enhancer activity and transcriptomic programs in undifferentiated ESCs^9^. These findings motivated us to study the roles of MLL3/4 enzymatic activities and MLL3/4-mediated enhancer H3K4me1 during development and cell differentiation. Using MLL3/4 enzyme-dead single and double knockin mice generated by CRISPR/Cas9, we found that MLL3/4 enzymatic activities are redundant and together are essential for gastrulation and early embryonic development. By knocking-in an MLL4 enzyme-dead point mutation in *Mll3*^-/-^ ESCs, we observed that the removal of MLL3/4 enzymatic activities does not impair the capacity of ESCs to differentiate towards all three germ layers, but leads to defects during the differentiation towards extraembryonic lineages. We attribute the failure in ExEn differentiation to impaired enhancer-binding of the lineage-determining TF GATA6. Consistent with cell culture data, *Sox2-Cre*-mediated elimination of MLL3/4 enzymatic activities in epiblast, but not in extraembryonic tissues, has little effect on gastrulation. Finally, we demonstrate that MLL3/4-catalyzed H3K4me1 is largely dispensable for enhancer activation during cell fate transition using EB differentiation and neural differentiation models.

## Results

### Loss of MLL3/4 enzymatic activities results in gastrulation failure and early embryonic lethality

MLL3 Tyr4792 (Y4792) and MLL4 Tyr5477 (Y5477) located in the catalytic SET domain are conserved among histone methyltransferases and are essential for enzymatic activity (Fig. 1a; Fig. 1-S1a)^9^. To investigate the role of MLL3/4 enzymatic activities in mouse development, we introduced the enzyme-dead point mutations Y4792A and Y5477A to *Mll3* and *Mll4* gene loci, respectively, by injecting CRISPR/Cas9 components into wild type zygotes (Fig. 1-S1b). After germline transmission, mice heterozygous for *Mll3* KI and *Mll4* KI (*Mll3*^KI/+^ and *Mll4*^KI/+^) survived without any discernible phenotypes and were inbred to obtain homozygotes. While *Mll3*^-/-^ mice display perinatal lethality^3, 7^, *Mll3*^KI/KI^ mice survived to adulthood with slightly reduced numbers and were fertile (Fig. 1b; Supplementary Table 1). *Mll4*^-/-^ mice die around embryonic day (E) 9.5^3, 7^. In contrast, *Mll4*^KI/KI^ mice died around birth with defective saccular structures in the lung (Fig. 1c; Fig. 1-S1c; Supplementary Table 1). These results suggest that MLL3 and MLL4 have both enzymatic activity-dependent and -independent functions in mouse development.

**Fig. 1:**
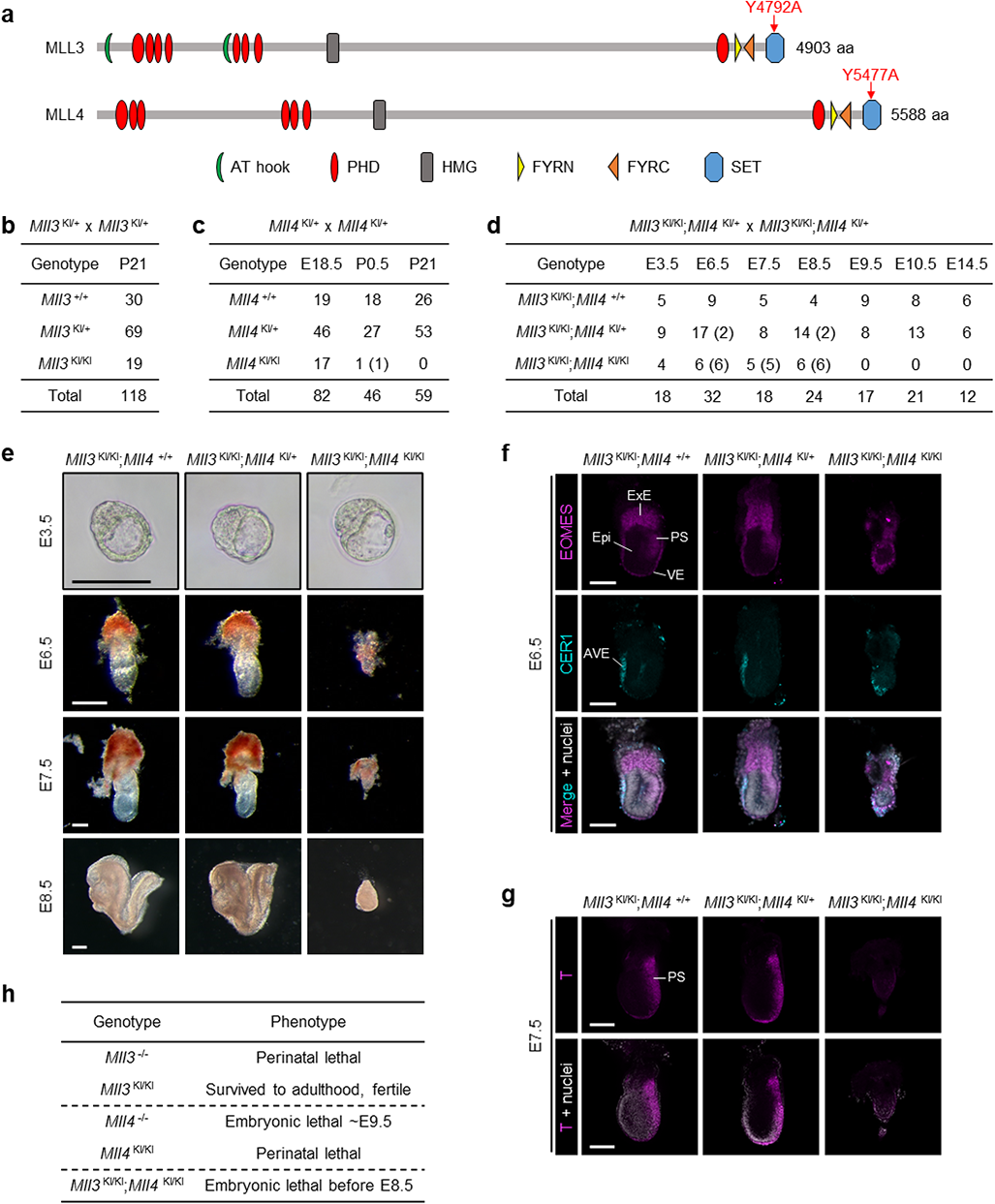
Loss of MLL3/4 enzymatic activities results in gastrulation failure and early embryonic lethality. **a**, Diagram of mouse MLL3 and MLL4 proteins. To eliminate the methyltransferase activities, tyrosine (Y) residues (highlighted in red) are mutated to alanine (A). PHD: Plant Homeotic Domain, HMG: High Mobility Group, FYRN/C: FY-Rich N/C-terminal, SET: Su(var)3-9, Enhancer-of-zeste and Trithorax. **b**, Genotypes of progeny from *Mll3*^KI/+^ intercrosses at postnatal day 21 (P21). **c**, Genotypes of progeny from *Mll4*^KI/+^ intercrosses at embryonic day 18.5 (E18.5), P0.5 and P21 are shown. Numbers of dead pups are indicated in parentheses. **d**, Genotypes of progeny from *Mll3*^KI/KI^;*Mll4*^KI/+^ intercrosses at indicated gestation stages are shown. Numbers of developmentally impaired embryos (E6.5 and E7.5) or embryonic remnants (E8.5) are indicated in parentheses. **e**, Representative images of E3.5 blastocysts and E6.5 - E8.5 embryos with indicated genotypes. Scale bar, 200 μm. **f**, Representative images of EOMES and CER1 immunofluorescence (IF) staining of E6.5 embryos. Scale bar, 100 μm. **g**, Representative images of T/BRACHYURY IF staining of E7.5 embryos. Scale bar, 200 μm. Cell nuclei were counter stained with DAPI (grey) in **f** and **g**. AVE, anterior visceral endoderm; Epi, epiblast; ExE, extraembryonic ectoderm; PS, primitive streak; VE, visceral endoderm. **h**, Summary of phenotypes of *Mll3*^-/-^, *Mll3*^KI/KI^, *Mll4*^-/-^, *Mll4*^KI/KI^, and *Mll3*^KI/KI^;*Mll4*^KI/KI^ mice. Phenotypes of *Mll3*^-/-^ and *Mll4*^-/-^ mice have previously been reported^3^.

Next, we crossed *Mll3* KI and *Mll4* KI mice to inactivate both MLL3 and MLL4. *Mll3*^KI/+^;*Mll4*^KI/+^ and *Mll3*^KI/KI^;*Mll4*^KI/+^ mice survived to adulthood and were fertile. However, no homozygous *Mll3*^KI/KI^;*Mll4*^KI/KI^ (double KI, dKI) embryos were recovered at E14.5 by intercrossing *Mll3*^KI/+^;*Mll4*^KI/+^ (Fig. 1-S1d; Supplementary Table 1). To determine at which stage dKI embryos could be detected, progeny from *Mll3*^KI/KI^;*Mll4*^KI/+^ intercrosses were examined. We failed to recover any dKI embryos beyond E8.5, suggesting they are early embryonic lethal (Fig. 1d; Fig. 1-S1e; Supplementary Table 1). At the blastocyst stage E3.5, dKI embryos were morphologically indistinguishable from control littermates (Fig. 1e). Immunofluorescence staining also revealed normal lineage allocation of epiblast, primitive endoderm and trophectoderm, marked by NANOG, GATA6 and CDX2, respectively, in E3.5 dKI blastocysts (Fig. 1-S1f). However, dKI embryos displayed obvious developmental defects at E6.5 and E7.5, and were only found as partially resorbed embryonic remnants at E8.5 (Fig. 1e).

We further analyzed E6.5 and E7.5 embryos by immunofluorescence staining. At E6.5, EOMES-labeling visceral endoderm (VE) underwent columnar-to-squamous transition and flattened out at the distal end in *Mll3*^KI/KI^;*Mll4*^+/+^ and *Mll3*^KI/KI^;*Mll4*^KI/+^ embryos as expected^7, 18, 19^, while the VE of dKI embryos was substantially thicker, suggesting a columnar-to-squamous transition defect (Fig. 1f). The directional migration of anterior visceral endoderm (AVE) during E5.5-6.5 is required for patterning the epiblast and to initiate gastrulation^20^. CER1, which labels the AVE^21^, was misexpressed in the distal region of dKI embryos at E6.5, suggesting an AVE migration failure (Fig. 1f). These observations indicate that VE development is profoundly compromised in dKI embryos. The onset of gastrulation is marked by the formation of primitive streak (PS), the precursor of mesoderm and definitive endoderm, in the posterior epiblast (Fig. 1-S1e)^12^. In *Mll3*^KI/KI^;*Mll4*^+/+^ and *Mll3*^KI/KI^;*Mll4*^KI/+^ embryos, we detected EOMES expression at E6.5 followed by abundant T expression at E7.5 in the PS as expected^11, 18^. In contrast, no specific EOMES or T signal was observed inside the epiblast of dKI embryos (Fig. 1f,g). Together, these data indicate that MLL3 and MLL4 enzymatic activities are largely redundant during embryonic development and that simultaneous elimination of both results in gastrulation failure and early embryonic lethality before E8.5 (Fig. 1h).

### ESCs lacking MLL3/4 enzymatic activities grow in monolayer but maintain cell identity

To investigate the function of MLL3/4 enzymatic activities *in vitro*, we took advantage of our previously derived *Mll3*^-/-^;*Mll4*^f/f^ (f/f) ESCs to eliminate the potential redundancy between MLL3 and MLL4^4^. Using CRISPR/Cas9 gene editing, we knocked-in the enzyme-dead point mutation Y5477A to both alleles of *Mll4* in f/f cells and generated independent *Mll3*^-/-^;*Mll4*^KI/KI^ (KI) ES cell lines (Fig. 2-S1a-c). We also re-generated *Mll3*/*Mll4* double knockout (KO) ESCs by transfecting a Cre-expressing plasmid into f/f ESCs^4^. The *Mll4* Y5477A point mutation in KI cell lines was confirmed by sequencing (Fig. 2-S1d). Further validation of KI cells found no evidence of potential exonic off-target mutations (Fig. 2-S1e-h). MLL4 protein was expressed at comparable levels in f/f and KI cells but absent in KO cells, and the MLL4-associated protein UTX followed a similar pattern (Fig. 2a)^3^. As expected, similarly reduced levels of global H3K4me1 and H3K4me2, but not H3K4me3, were observed in KI and KO ESCs (Fig. 2b).

**Fig. 2:**
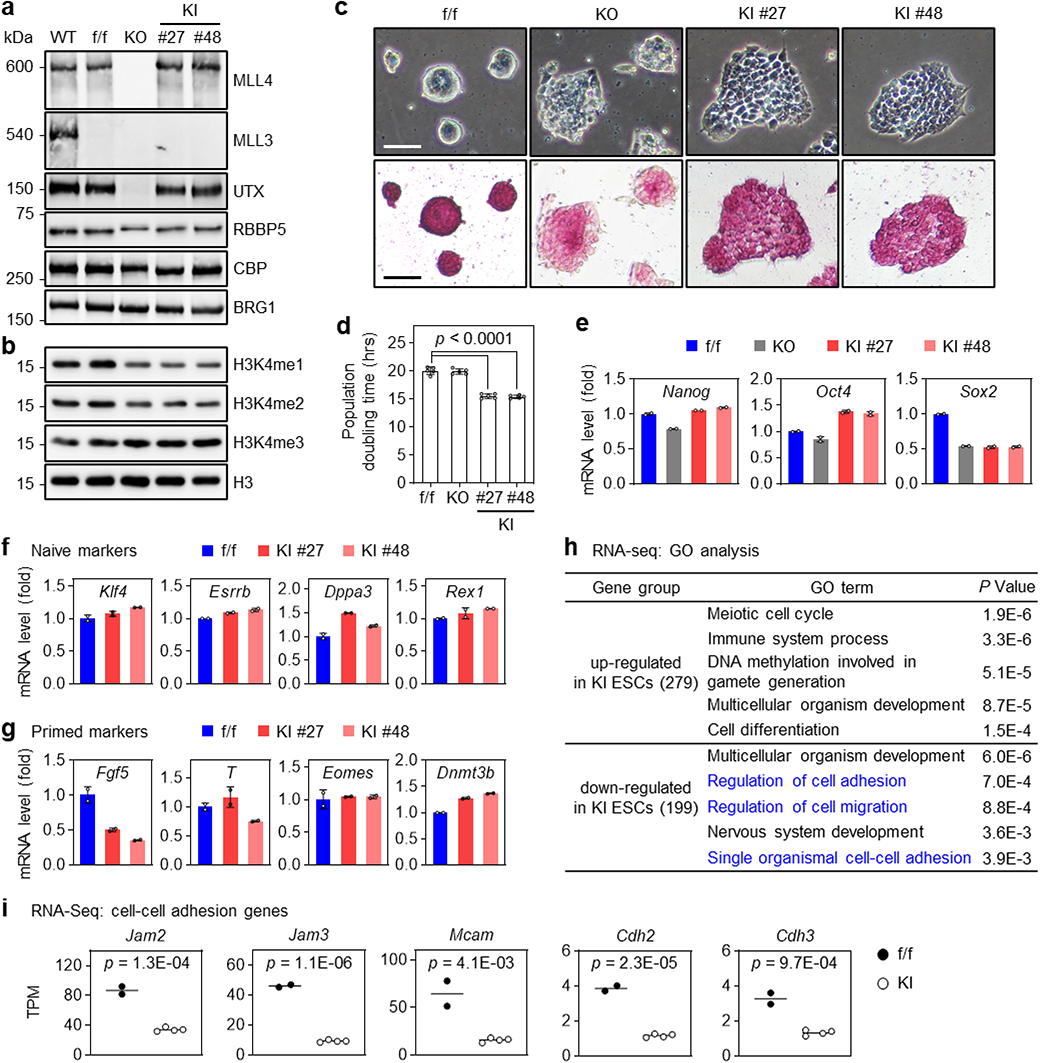
ESCs lacking MLL3/4 enzymatic activities grow in monolayer but maintain cell identity. **a**,**b**, MLL3/4 enzyme-dead (*Mll3*^-/-^;*Mll4*^KI/KI^, KI) ESCs were generated as described in Fig. 2-S1. Nuclear extracts (**a**) or histone extracts (**b**) prepared from wild type (WT), *Mll3*^-/-^;*Mll4*^f/f^ (f/f), *Mll3*/*Mll4* double knockout (KO) and KI ESCs were analyzed with immunoblotting using indicated antibodies. **c**, Representative phase contrast microscopic (*upper*) and AP staining (*lower*) images. Scale bar, 50 μm. **d**, Population doubling time. Data are presented as means ± SD (*n* = 5). Statistical significance was determined by the two-tailed unpaired *t*-test. **e**, Pluripotency markers were analyzed by RT-qPCR (*n* = 2). **f**,**g**, ESCs lacking MLL3/4 enzymatic activities maintain ES cell identity. Naïve markers (**f**) and primed markers (**g**) were analyzed by RT-qPCR (*n* = 2). **h**, GO analysis of up-regulated and down-regulated genes in KI ESCs. The number of genes in each group is indicated in parentheses. Terms associated with cell adhesion are highlighted in blue. **i**, Expression levels of representative genes associated with cell-cell adhesion. Data from RNA-seq are presented as dot plots (f/f, n = 2; KI, n = 4). Horizontal lines represent mean values. Statistical significance was determined by the two-tailed unpaired *t*-test.

Next, we characterized the morphology and cell growth of KI ESCs. When cultured in 2i+LIF medium, neither KO nor KI ESCs could form regular dome-like colonies shown in f/f ESCs. KI colonies were flat, and cells grew homogeneously in monolayer (Fig. 2c). Interestingly, KI ESCs grew faster and displayed ∼20% decrease in population doubling time (Fig. 2d). Expression levels of pluripotency markers *Nanog* and *Oct4* were largely comparable among all cell lines, except that *Sox2* levels decreased about two-fold in both KI and KO ESCs as reported previously (Fig. 2e)^22^. In addition, both f/f and KI ESCs presented positive alkaline phosphatase staining (Fig. 2c).

ESCs maintain naïve pluripotency while epiblast stem cells retain primed pluripotency and show flat morphology^23^. Although KI ESCs also showed flat morphology, expression levels of naïve and primed markers were largely comparable between f/f and KI ESCs (Fig. 2f,g). Interestingly, by RNA-seq analysis we found that genes down-regulated in KI ESCs were enriched for terms related to cell adhesion (Fig. 2h). Indeed, lower expression of representative cell-cell adhesion genes was observed in KI ESCs (Fig. 2i). Together, these data indicate that while KI ESCs show monolayer growth, possibly due to impaired cell-cell adhesion, MLL3/4 enzymatic activities are dispensable for maintaining cell identity.

### ESCs lacking MLL3/4 enzymatic activities can differentiate towards the three germ layers

We next investigated the role of MLL3/4 enzymatic activities in EB differentiation. During EB differentiation, MLL4 was expressed at similar levels between f/f and KI cells, indicating that loss of enzymatic activity did not affect MLL4 protein stability (Fig. 3-S1a). As expected, global H3K4me1 and H3K4me2 decreased in KI EBs (Fig. 3-S1b). Consistent with our previous report^4^, f/f ESCs formed large cystic EBs upon differentiation whereas KO cells displayed severe defects (Fig. 3a,b). Although none of KI EBs displayed cystic cavities, they exhibited similar sizes as f/f EBs throughout differentiation (Fig. 3a,b). After differentiation, both f/f and KI cells showed dramatically decreased expression of pluripotency markers and increased expression of markers for the three germ layers (Fig. 3c; Fig. 3-S1c). In addition, PS markers were temporally expressed at similar levels in f/f and KI D4 EBs (Fig. 3-S1a,c).

**Fig. 3:**
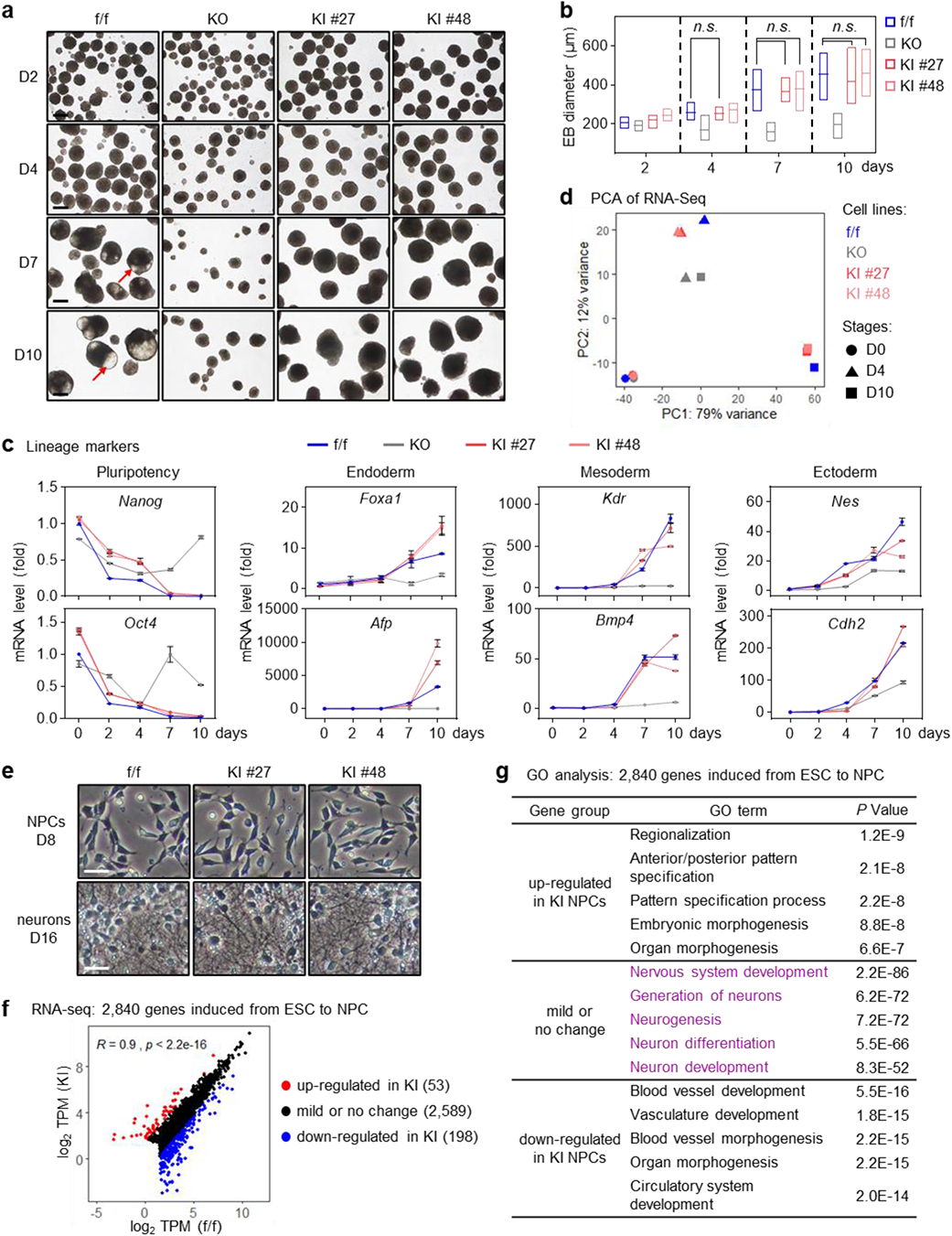
ESCs lacking MLL3/4 enzymatic activities can differentiate towards the three germ layers. **a-d**, Embryoid body (EB) differentiation from f/f, KO and KI ESCs. **a**, Representative microscopic images at day 2 (D2), D4, D7 and D10. Representative cystic cavities are indicated with red arrows. Scale bar, 250 μm. **b**, Diameters of EBs at indicated time points are presented as interleaved low-high floating bar plots. Center lines represent mean values. At each time point, at least 17 EBs were measured per group. Statistical significance was determined by the two-tailed unpaired *t*-test. **c**, RT-qPCR analysis of pluripotency markers (*Nanog*, *Oct4*), endoderm markers (*Foxa1*, *Afp*), mesoderm markers (*Kdr*, *Bmp4*) and ectoderm markers (*Nes*, *Cdh2*) at indicated time points. **d**, Principal component analysis (PCA) of RNA-seq data at D0, D4 and D10. **e-g**, f/f and KI ESCs were induced to differentiate into neural progenitor cells (NPCs) and neurons sequentially. D8 NPCs and D16 neurons were collected for RNA-seq. **e**, Representative microscopic images of D8 NPCs (*upper*) and D16 neurons (*lower*). Scale bar, 25 μm. **f**, Expression levels of 2,840 genes induced from ESC to NPC were presented as scatter plots. Pearson correlation coefficient (*R*) and *P* value are shown. The number of genes in each group is indicated in parentheses. **g**, GO analysis of three gene groups in **f**. Terms associated with neural differentiation are highlighted in purple.

We then performed RNA-seq analysis during EB differentiation at D0, D4 and D10. Principal component analysis (PCA) showed that transcriptomes of KI cells but not KO cells resembled those of f/f cells (Fig. 3d). For most developmental processes, gene set enrichment analysis (GSEA) did not reveal significant differences in gene expression between f/f and KI EBs at D10, when most markers for the three germ layers are strongly induced. One exception is cardiac muscle development, a term significantly associated with genes down-regulated in KI EBs (Fig. 3-S1d). Consistently, we observed delayed formation of spontaneous beating clusters, decreased beating frequency and compromised induction of cardiomyogenesis related genes during EB differentiation of KI ESCs (Fig. 3-S1e-h). To validate the *in vivo* differentiation capacity of KI ESCs, we performed a teratoma assay. Both f/f and KI ESCs developed to teratomas containing tissues derived from all three germ layers (Fig. 3-S1i). Together, these data indicate that despite defects in EB cavitation and cardiomyogenesis, KI ESCs are capable of differentiating towards the three germ layers.

We also examined the role of MLL3/4 enzymatic activities in germ-layer specification using neural differentiation as a model (Fig. 3-S2a)^16^. We observed that both f/f and KI ESCs differentiated homogeneously to neural progenitor cells (NPCs) around D8 with characteristics of spindle-like shape and neural rosette; after D10, NPCs gradually differentiated to mature glutamatergic neurons until D16, when a dense neuritic network formed (Fig. 3e; Fig. 3-S2b). In contrast, KO cells showed severe differentiation defects even at the NPC stage (Fig. 3-S2b). RNA-seq analyses in NPCs and neurons revealed that only those genes induced to comparable levels between f/f and KI cells were functionally associated with neural lineage commitment (Fig. 3f,g; Fig. 3-S2c,d). Consistently, NPC and neuron markers were expressed at similar levels in f/f and KI cells (Fig. 3-S2e,f). These results indicate that while MLL3/4 proteins are essential for neural differentiation, their enzymatic activities are generally dispensable.

### Loss of MLL3/4 enzymatic activities impairs ExEn gene induction during EB differentiation

D3-D5 EBs contain cell lineages found in E6.5-7.0 embryos during gastrulation, such as an outer layer ExEn and the transiently formed PS-like cells^15^. Because dKI embryos showed severe developmental defects at E6.5 (Fig. 1e-g), we performed GSEA at D4 EB using cell-type markers defined previously by single-cell RNA-seq in E6.5 embryos^24^. Interestingly, markers of VE and parietal endoderm, the two ExEn lineages, were significantly enriched in genes down-regulated in KI EBs, while markers of PS or nascent mesoderm, an early derivative of PS, showed no enrichment (Fig. 4-S1a; Fig. 1-S1e).

We also compared the expression of reported cell-type markers^12^ between f/f and KI D4 EBs. All 12 ExEn markers tested were expressed at lower levels in KI EBs, with 10 of them showing over 2-fold decrease. In contrast, no PS markers changed more than 2-fold (Fig. 4-S1b). BMP, Nodal and Wnt signaling pathways play key roles in early embryonic development^12^. Among all examined components of these pathways, only *Bmp2* and *Bmp4*, which encode two ligands involved in VE development^25^, decreased more than 2-fold in KI EBs (Fig. 4-S1b). We confirmed the reduced expression of ExEn genes in KI D4 EBs by RT-qPCR analysis and immunofluorescence staining (Fig. 4-S1c,d). Once EBs attach to gelatinized surfaces, ExEn cells migrate away from the EB periphery and form a ring of epithelial cells^26^. Consistent with the reduced ExEn gene expression, we noticed that fewer cells dispersed and migrated out of attached KI EBs compared to f/f EBs (Fig. 4-S1e). Together, these data indicate that MLL3/4 enzymatic activities are required for induction of ExEn genes during EB differentiation.

### ESCs lacking MLL3/4 enzymatic activities show aberrant differentiation to extraembryonic lineages

Next, we investigated whether MLL3/4 enzymatic activities are required for ExEn differentiation using a published protocol (Fig. 4a). Five days after induction of differentiation, we observed well-developed ExEn colonies formed from f/f ESCs with characteristics of cobblestone-like morphology. In contrast, KI ESCs failed to form any ExEn colonies (Fig. 4b). GATA6 is the lineage-determining TF at the top of the regulatory hierarchy governing ExEn lineage commitment^27^. We found that at both mRNA and protein levels, *Gata6* expression was comparable between f/f and KI cells after ExEn differentiation, while the induction of down-stream ExEn markers, such as *Gata4* and *Sox17*, was severely impaired in KI cells (Fig. 4c,d; Fig. 4-S2a). These data indicate that loss of MLL3/4 enzymatic activities in ESCs leads to severe defects in ExEn differentiation and GATA6-dependent transcriptional program.

**Fig. 4:**
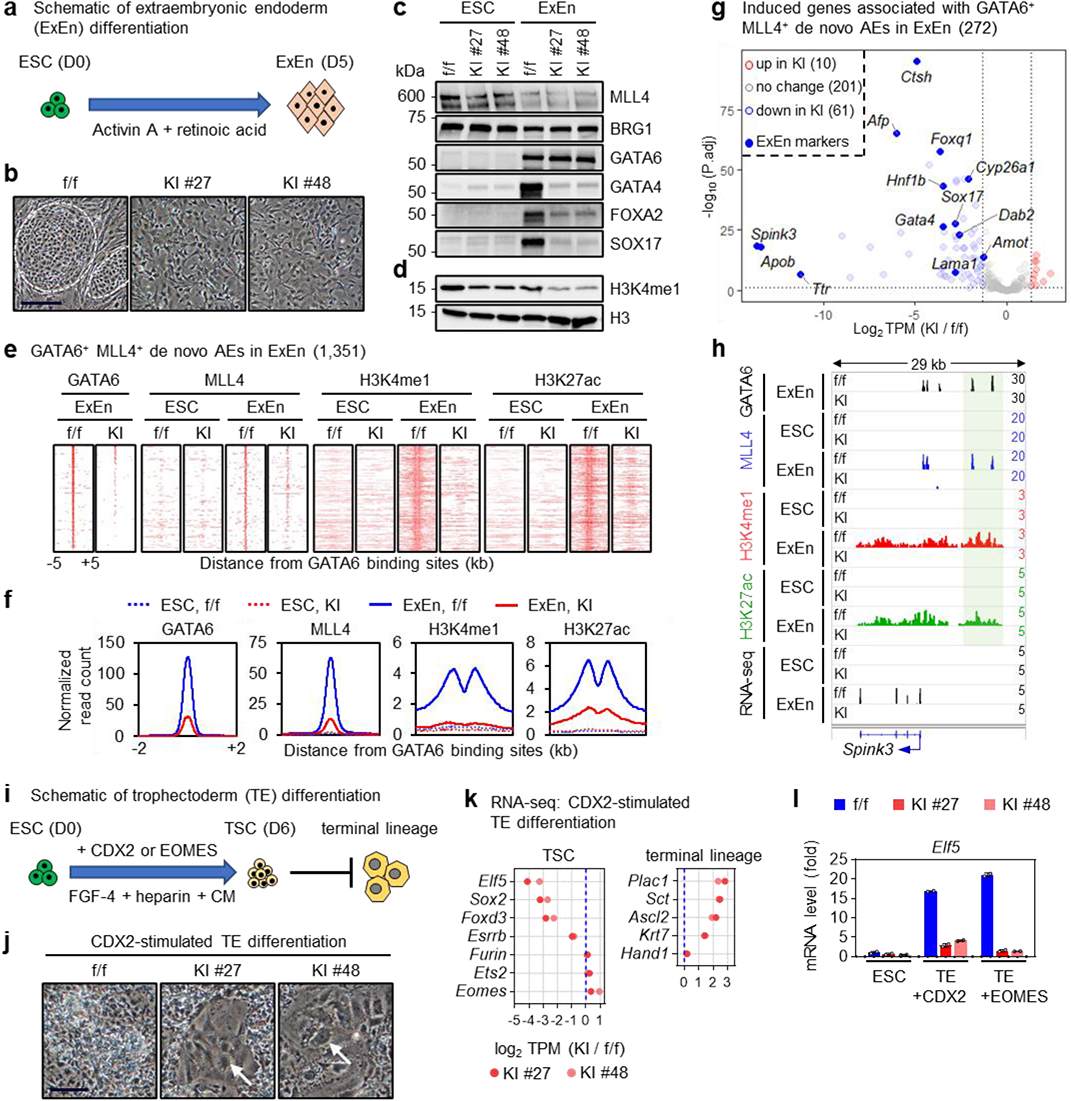
ESCs lacking MLL3/4 enzymatic activities show aberrant differentiation to extraembryonic lineages. **a**, Schematic of extraembryonic endoderm (ExEn) differentiation. After 5-day differentiation, ESCs are converted to ExEn cells. **b**, Representative microscopic images of cells after ExEn differentiation. Representative ExEn colonies are indicated with white circles. Scale bar, 200 μm. **c**,**d**, Immunoblotting in f/f and KI cells before and after ExEn differentiation. Whole cell lysates (**c**) or histone extracts (**d**) were analyzed using indicated antibodies. **e-h**, After ExEn differentiation, cells were collected for ChIP-seq and RNA-seq. Heat maps (**e**) and average profiles (**f**) of GATA6 and MLL4 genomic bindings as well as H3K4me1 and H3K27ac enrichments on GATA6^+^ MLL4^+^ de novo active enhancers (AEs) in ExEn cells are shown. **g**, Expression levels of 272 induced genes associated with GATA6^+^ MLL4^+^ de novo AEs in ExEn cells were presented as volcano plots. The number of genes in each group is indicated in parentheses. ExEn markers are highlighted as blue dots. **h**, ChIP-seq profiles of GATA6, MLL4, H3K4me1 and H3K27ac as well as RNA-seq profiles in f/f and KI cells are displayed on the *Spink3* locus. GATA6^+^ MLL4^+^ de novo AEs are highlighted in shades. **i**, Schematic of trophectoderm (TE) differentiation. After 6-day differentiation, ESCs are converted to trophoblast stem cells (TSCs). The culture condition supports maintenance of TSCs and prevents terminal trophoblast differentiation. **j**, Representative microscopic images of cells after CDX2-stimulated TE differentiation. Representative cells belonging to terminal lineages are indicated with white arrows. Scale bar, 100 μm. **k**, Expression fold changes of TSC markers and terminal trophoblast lineage markers between KI and f/f cells after CDX2-stimulated TE differentiation. **l**, RT-qPCR analysis of *Elf5* expression before and after CDX2-or EOMES-stimulated TE differentiation (*n* = 2).

To uncover the underlying mechanism, we conducted ChIP-seq of GATA6, MLL4, H3K4me1 and H3K27ac. Strikingly, we observed that the number of GATA6-bound active enhancers (AEs) decreased dramatically in KI cells compared to f/f cells after ExEn differentiation (Fig. 4-S2b). Reduced enhancer-binding of GATA6 led to diminished recruitment of MLL4 and the failure in enhancer activation as indicated by H3K27ac induction (Fig. 4e,f). The physical interaction between MLL4 and GATA6 was not affected in KI cells (Fig. 4-S2c). RNA-seq analysis of genes induced after differentiation and associated with GATA6^+^ MLL4^+^ de novo AEs revealed that 22% of them were down-regulated in KI cells and were functionally associated with endoderm development (Fig. 4g,h; Fig. 4-S2d,e).

We also examined the role of MLL3/4 enzymatic activities in ESC differentiation to trophectoderm, the other extraembryonic lineage (Fig. 1-S1e). By overexpressing a trophectoderm lineage-determining TF, such as CDX2 or EOMES, ESCs can be converted to and maintained as trophoblast stem cells (TSCs) (Fig. 4i; Fig. 4-S2g)^28^. Six days after CDX2-stimulated trophectoderm differentiation, flattened TSC colonies formed from f/f ESCs as expected. Surprisingly, KI ESCs showed precocious trophectoderm differentiation and developed to numerous giant cells, which are terminally differentiated trophoblast (Fig. 4j)^29^. Transcriptomic analysis revealed decreased expression of TSC markers and increased expression of terminal trophoblast lineage markers in KI cells compared to f/f cells after trophectoderm differentiation (Fig. 4k). Among TSC markers down-regulated in KI cells, *Elf5* was the most prominent one (Fig. 4k). The induction of *Elf5* was also blocked in EOMES-stimulated trophectoderm differentiation of KI ESCs (Fig. 4l). ELF5 is a key TF governing TSC identity and is necessary to prevent terminal trophoblast differentiation^29^. Our observations indicate that loss of MLL3/4 enzymatic activities blocks induction of *Elf5* during ESC to trophectoderm transition and causes precocious terminal trophoblast differentiation.

### MLL3/4 enzymatic activities are required for GATA6 binding on enhancers

GATA6 overexpression is sufficient to reprogram ESCs into ExEn cells in the presence of pluripotency signals^30^. To directly evaluate the effect of MLL3/4 enzymatic activity loss on GATA6 genomic binding and GATA6-driven ESC reprogramming, we ectopically expressed GATA6 in f/f and KI ESCs using a doxycycline (Dox)-inducible system (Fig. 5a,b). ChIP-seq analysis performed 1 day after GATA6 induction by Dox treatment revealed that GATA6 binding on enhancers was lost in KI cells (Fig. 5c,d). Recruitment of MLL4 and the consequent enhancer activation were also impaired on GATA6 induced AEs (Fig. 5d). After 3 days of Dox treatment, we observed the formation of cobblestone-like ExEn colonies as well as robust induction of GATA6 target genes *Gata4* and *Sox17* in f/f cells but not in KI cells (Fig. 5e,f).

**Fig. 5:**
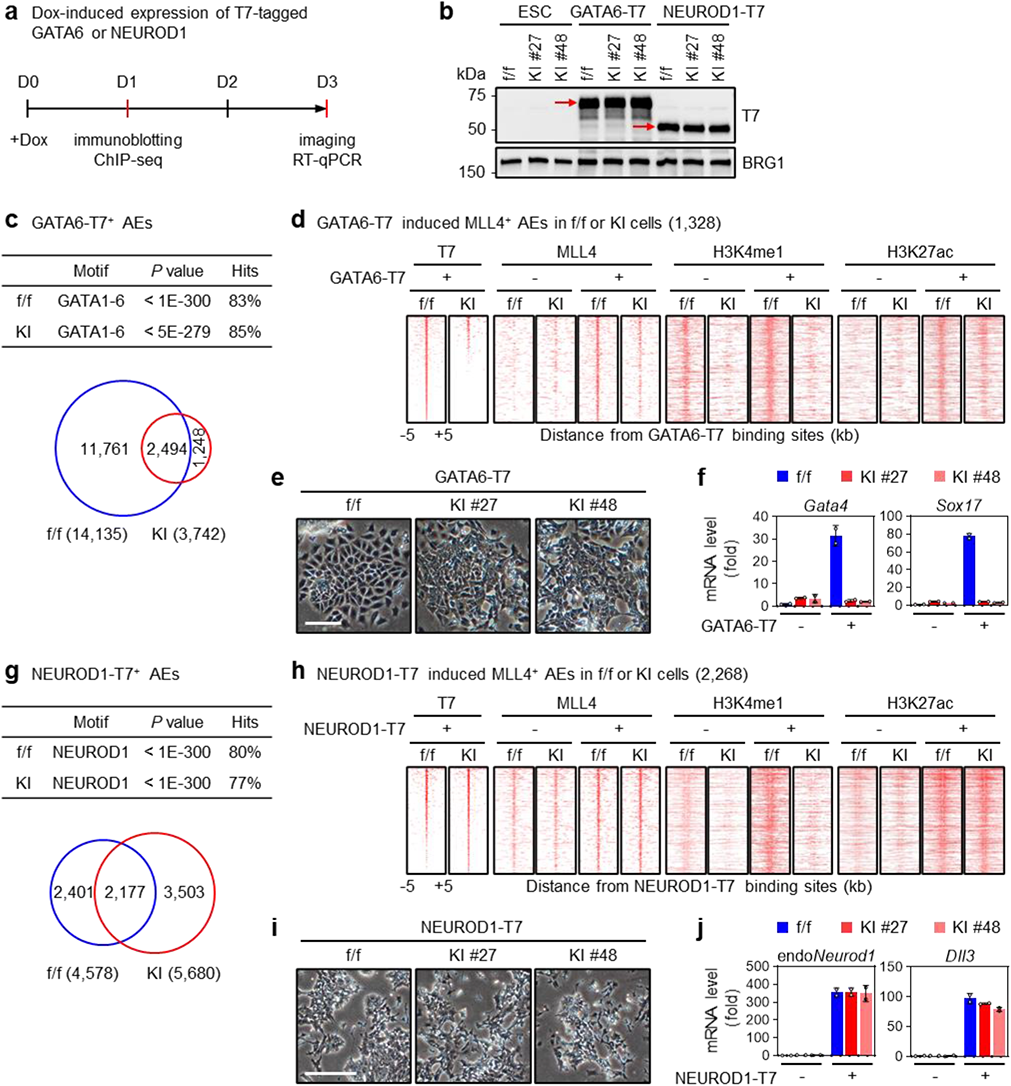
MLL3/4 enzymatic activities are required for GATA6 binding on enhancers. **a**, Schematic of Dox-induced GATA6 or NEUROD1 expression. f/f and KI ESCs were infected with Doxycycline (Dox)-inducible lentiviral vector expressing T7-tagged GATA6 or NEUROD1. Cells were treated with 1 μg/ml Dox. Immunoblotting and ChIP-seq were performed at D1, imaging and RT-qPCR were performed at D3. **b**, Immunoblotting of T7-tagged GATA6 and NEUROD1. Whole cell lysates were analyzed using indicated antibodies. BRG1 is shown as loading control. **c**, Motif analysis and Venn diagram of GATA6^+^ AEs in f/f and KI cells. **d**, Heat maps of GATA6-T7 and MLL4 genomic bindings as well as H3K4me1 and H3K27ac enrichments on GATA6-T7^+^ MLL4^+^ AEs induced by GATA6-T7 overexpression. **e**, Representative microscopic images of cells after Dox-induced expression of GATA6-T7. Scale bar, 100 μm. **f**, GATA6 target genes were analyzed by RT-qPCR before and after Dox-induced expression of GATA6-T7 (*n* = 2). **g**, Motif analysis and Venn diagram of NEUROD1^+^ AEs in f/f and KI cells. **h**, Heat maps of NEUROD1-T7 and MLL4 genomic bindings as well as H3K4me1 and H3K27ac enrichments on NEUROD1-T7^+^ MLL4^+^ AEs induced by NEUROD1-T7 overexpression. **i**, Representative microscopic images of cells after Dox-induced expression of NEUROD1-T7. Scale bar, 200 μm. **j**, NEUROD1 target genes were analyzed by RT-qPCR before and after Dox-induced expression of NEUROD1-T7 (*n* = 2).

As a comparison, we did similar analyses in ESCs ectopically expressing NEUROD1, a lineage-determining TF capable of reprogramming ESCs towards the neural lineage (Fig. 5a,b)^31^. Different from the impact on GATA6, loss of MLL3/4 enzymatic activities did not reduce the number of NEUROD1-bound AEs (Fig. 5g,h). NEUROD1 recruited MLL4 and induced enhancer activation to a similar extent in f/f and KI cells (Fig. 5h). Moreover, within 3 days of NEUROD1 induction, both f/f and KI cells exhibited a spindle-like NPC morphology and expressed NEUROD1 target genes, indicating successful reprogramming (Fig. 5i,j). Together, these data suggest that MLL3/4 enzymatic activities are selectively required for GATA6 enhancer-binding and GATA6-driven ESC reprogramming to the ExEn lineage.

### Lineage-selective roles of MLL3/4 enzymatic activities in early embryonic development

KI ESCs can differentiate towards the three germ layers but show aberrant differentiation towards extraembryonic lineages, implying that loss of MLL3/4 enzymatic activities in extraembryonic, rather than embryonic, tissues is responsible for the gastrulation failure and early embryonic lethality of dKI mice. Therefore, we took a conditional KI approach and selectively eliminated MLL3/4 enzymatic activities in embryonic tissues using the epiblast-specific *Sox2-Cre* transgene (Fig. 1-S1e)^32^. Among progeny from crossing *Mll3*^KI/KI^;*Mll4*^KI/+^;*Sox2-Cre* male with *Mll3*^f/f^;*Mll4SET*^f/f^ female mice^10^, 25% are expected to be *Sox2-Cre*-driven conditional MLL3/4 double KI embryos (*Mll3* ^f/KI^;*Mll4SET* ^f/KI^;*Sox2-Cre*, referred to as Sox2-dKI hereafter) and 25% be controls (*Mll3* ^f/KI^;*Mll4SET* ^f/+^;*Sox2-Cre*, referred to as Sox2-Ctr hereafter) (Fig. 6-S1a,b). In marked contrast to constitutive dKI embryos (Fig. 1d), Sox2-dKI embryos were recovered at the expected Mendelian ratio up to E12.5 (Fig. 6a; Supplementary Table 1). At E7.5 and E8.5, there were no noticeable morphological differences between Sox2-dKI and Sox2-Ctr embryos. At E9.5 and E10.5, Sox2-dKI embryos were still viable but displayed developmental abnormalities including failed embryonic turning and pericardial edema. At E12.5, Sox2-dKI embryos underwent resorption (Fig. 6b). *Sox2-Cre*-mediated elimination of MLL3/4 enzymatic activities was confirmed by immunoblotting of H3K4me1 in E8.5 and E9.5 embryos (Fig. 6c; Fig. 6-S1c).

**Fig. 6:**
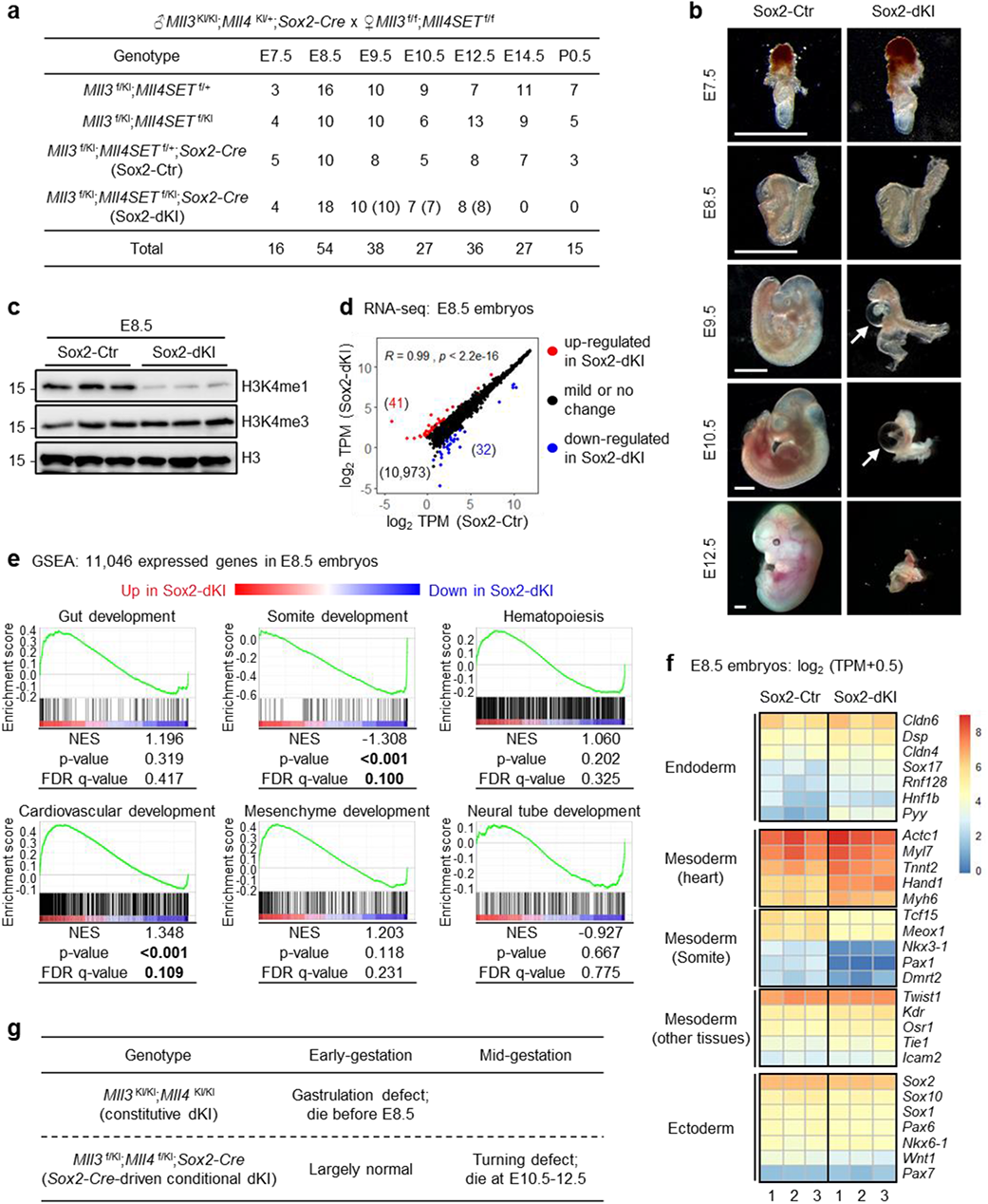
Lineage-selective roles of MLL3/4 enzymatic activities in early embryonic development. **a**, Genotypes of progeny from mating between *Mll3*^KI/KI^;*Mll4*^KI/+^;*Sox2-Cre* male and *Mll3*^f/f^;*Mll4SET*^f/f^ female mice at indicated gestation stages. Numbers of developmentally impaired embryos (E9.5 and E10.5) or embryonic remnants (E12.5) are indicated in parentheses. **b**, Representative images of E7.5 - E10.5 and E12.5 embryos with indicated genotypes. White arrows indicate pericardial edema. Scale bar, 1 mm. **c**, Whole cell lysates prepared from E8.5 yolk sac-removed embryos were analyzed with immunoblotting using indicated antibodies. **d**, Sox2-Ctr and Sox2-dKI embryos were collected at E8.5 for RNA-seq. Levels of expressed genes were presented as scatter plots. Pearson correlation coefficient (*R*) and *P* value are shown. The number of genes in each group is indicated in parentheses. **e**, GSEA of 11,046 expressed genes in E8.5 embryos on developmental terms. NES, normalized enrichment score. Statistically significant data are highlighted in bold. **f**, Heat maps of gene expression level in E8.5 embryos for markers of germ layer derivatives. Data from 3 embryos were presented for each genotype. **g**, Summary of developmental stage-specific phenotypes of constitutive double KI (dKI) and *Sox2-Cre*-driven conditional dKI mice.

RNA-seq analysis revealed that transcriptomic patterns were highly comparable between Sox2-dKI and Sox2-Ctr embryos at E8.5 and E9.5 (Fig. 6d; Fig. 6-S1d). For most key processes of early organogenesis, E8.5 or E9.5 Sox2-dKI embryos did not show significant gene expression changes compared to Sox2-Ctr embryos by GSEA, indicating the three germ layers are successfully developed during gastrulation of Sox2-dKI embryos (Fig. 6e; Fig. 6-S1e). Interestingly, the term “cardiovascular development” was enriched in up-regulated genes, while the term “somite development” was enriched in down-regulated genes of Sox2-dKI embryos, which may partially explain mid-gestational defects observed in Sox2-dKI embryos (Fig. 6e; Fig. 6-S1e). In agreement with the GSEA result, genes specific for tissues derived from the three germ layers^33^ were expressed at similar levels in Sox2-dKI and Sox2-Ctr embryos, although markers for heart and somite were differentially expressed (Fig. 6f; Fig. 6-S1f). Together, these data indicate that embryos lacking MLL3/4 enzymatic activities selectively in embryonic tissues can develop through gastrulation until mid-gestation, thus suggesting that the gastrulation failure observed in constitutive dKI embryos is due to loss of MLL3/4 enzymatic activities in extraembryonic tissues (Fig. 6g). Our data also suggest that MLL3/4 enzymatic activities play lineage-selective roles in early embryonic development.

### MLL3/4-catalyzed H3K4me1 is largely dispensable for enhancer activation during ESC differentiation

MLL3/4 proteins are required for enhancer activation during EB differentiation^4^. To characterize the role of MLL3/4 enzymatic activities in de novo enhancer activation, we conducted ChIP-seq of MLL4, H3K4me1 and H3K27ac in f/f and KI D4 EBs. We noticed that loss of enzymatic activity resulted in redistribution of MLL4 genomic occupancy without discernible changes of total binding intensity (Fig. 7-S1a). We identified 10,767 de novo MLL4^+^ AEs that were present in f/f or KI D4 EBs but not in ESCs, and divided them into three groups based on the comparison of MLL4 binding intensities between f/f and KI EBs: Group I (1,504, 14%), increased in KI EBs; Group II (3,610, 34%), comparable between f/f and KI EBs; Group III (5,653, 52%), decreased in KI EBs (Fig. 7-S1b). Interestingly, motifs of GATA family TFs including GATA6 were strongly enriched only in Group III AEs (Fig. 7-S1c). Diminished GATA6 binding was confirmed in KI EBs by ChIP-seq, which likely explains the impaired MLL4 genomic occupancy on Group III AEs (Fig. 7-S1d).

To test if the requirement of MLL3/4 proteins for de novo enhancer activation is mediated by MLL3/4-catalyzed H3K4me1, we examined distributions of H3K4me1, H3K27ac and chromatin accessibility, assessed by ATAC-seq, on Group II de novo MLL4^+^ AEs where f/f and KI EBs display similar levels of MLL4 binding. We observed dramatic induction of H3K4me1, H3K27ac and chromatin accessibility from ESC to D4 EB in f/f cells. In KI EBs, despite no gain of H3K4me1, Group II AEs showed comparable induction of H3K27ac and chromatin accessibility as in f/f EBs (Fig. 7a,b). Genes associated with Group II AEs were also expressed at similar levels between f/f and KI EBs (Fig. 7-S1e). Using ChIP-seq data from KO cells^4^, we further confirmed the requirement of MLL3/4 proteins for the activation of Group II AEs (Fig. 7-S2a,b). Representative genes associated with Group II AEs were induced comparably in f/f and KI but not KO EBs (Fig. 7c; Fig. 7-S2c). On the other hand, induction patterns of H3K27ac and chromatin accessibility on Group I and III AEs were consistent with those of MLL4 but not H3K4me1 (Fig. 7a,b). The reduced H3K27ac and chromatin accessibility on Group III AEs could be attributed to reduced MLL4 genomic occupancy, although we cannot exclude the contribution of the H3K4me1 loss. Expression levels of genes associated with Group I or Group III AEs were significantly up- or down-regulated in KI EBs compared to f/f EBs (Fig. 7-S1e). Together, these data indicate that while MLL3/4 proteins are required for enhancer activation during early EB differentiation, MLL3/4-catalyzed H3K4me1 is dispensable, at least on Group II AEs.

**Fig. 7:**
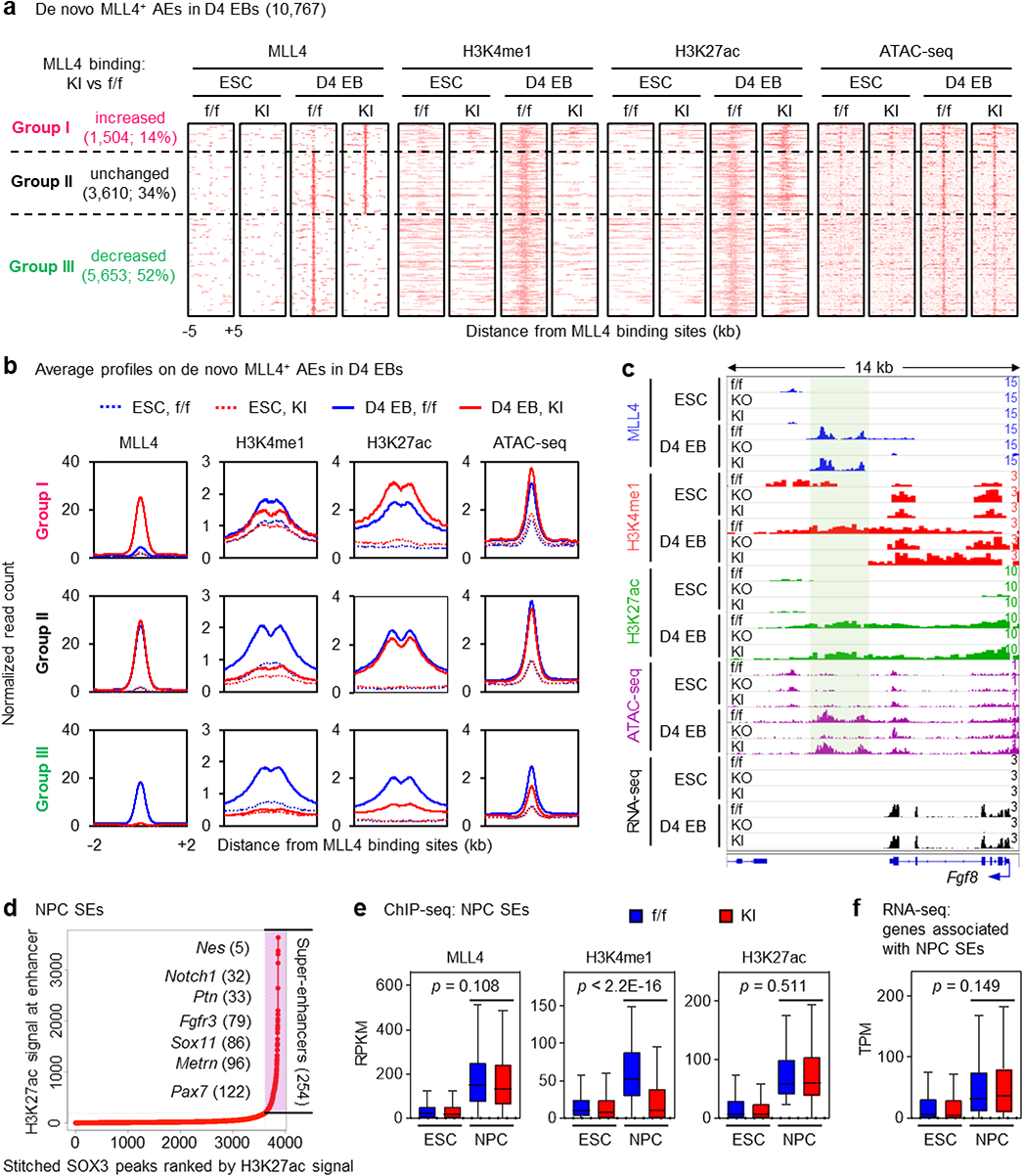
MLL3/4-catalyzed H3K4me1 is largely dispensable for enhancer activation during ESC differentiation. **a-c**, D4 EBs were collected for ChIP-seq and ATAC-seq. Heat maps (**a**) and average profiles (**b**) of MLL4 genomic bindings, H3K4me1 and H3K27ac enrichments as well as ATAC-seq signals on the three groups of de novo MLL4^+^ AEs identified in Fig. 7-S1b are shown. **c**, ChIP-seq profiles of MLL4, H3K4me1 and H3K27ac, ATAC-seq and RNA-seq profiles in f/f, KO and KI cells are displayed on the *Fgf8* locus. Group II de novo MLL4^+^ AE are highlighted in shades. ChIP-seq data in KO cells were obtained from GSE50534^4^. **d-f**, D8 NPCs were collected for ChIP-seq. **d**, NPC super-enhancers (SEs) were identified by stitching SOX3 binding sites in NPC AEs. SOX3 ChIP-seq data were obtained from GSE33059^35^. SEs were ranked by H3K27ac signal intensities. Representative genes associated with NPC SEs are indicated. The SE rank of each gene is indicated in parentheses. **e**, MLL4, H3K4me1 and H3K27ac intensities in ESCs and NPCs on NPC SEs are presented in box plots. **f**, Expression levels in ESCs and NPCs of genes associated with NPC SEs are presented in box plots. **e**,**f**, Center lines represent median values; the bottom and top of the boxes represent lower and upper quartiles; whiskers were calculated using the Tukey method. Statistical significance was determined by the two-sided Wilcoxon signed-rank test.

We also investigated the role of MLL3/4-catalyzed H3K4me1 in enhancer activation during neural differentiation. Super-enhancers (SEs) are clusters of AEs bound by lineage-specific master TFs and drive high-level expression of cell-identity genes^34^. SOX3 is the master TF in NPCs^35^. We identified 254 SEs in f/f NPCs using stitched SOX3 peaks^35^ ranked by H3K27ac signal intensity. 97.6% (248/254) of them were MLL4^+^ and associated with NPC identity genes such as *Nes*, *Notch1* and *Ptn* (Fig. 7d). GO analysis of SE-associated genes identified “nervous system development” as the top term (Fig. 7-S3a). While induction of H3K4me1 signals on NPC SEs displayed severe impairment in KI NPCs, signal intensities of MLL4 and H3K27ac did not show significant differences between f/f and KI NPCs. Expression levels of SE-associated genes were also induced comparably (Fig. 7e,f; Fig. 7-S3b). These results demonstrate that MLL3/4-catalyzed H3K4me1 is dispensable for SE activation during neural differentiation.

## Discussion

In this study, we show that MLL3/4 carry both enzymatic activity-dependent and -independent functions and that MLL3/4 enzymatic activities are redundant during early embryonic development in mice. This is the first *in vivo* evidence of functional redundancy between enzymatic activities of Set1-like H3K4 methyltransferases. Constitutive elimination of both MLL3/4 enzymatic activities leads to gastrulation failure while *Sox2-Cre*-mediated elimination of them in embryonic tissues leaves gastrulation largely intact. In culture, ESCs lacking MLL3/4 enzymatic activities can differentiate towards all three embryonic germ layers but show aberrant differentiation to extraembryonic lineages, which provides a potential explanation for the observed phenotypes in mice. Moreover, our data reveal that MLL3/4-catalyzed H3K4me1 is largely dispensable for enhancer activation in ESC differentiation. We also find that eliminating MLL3/4 enzymatic activities in ESCs impedes GATA6 binding on enhancers, which likely causes the failure in ExEn lineage commitment. Based on our data, we propose a model that MLL3/4 enzymatic activities regulate ESC differentiation and early embryonic development through selectively modulating genomic binding of lineage-determining TFs, rather than directly activating enhancers (Fig. 8).

**Fig. 8:**
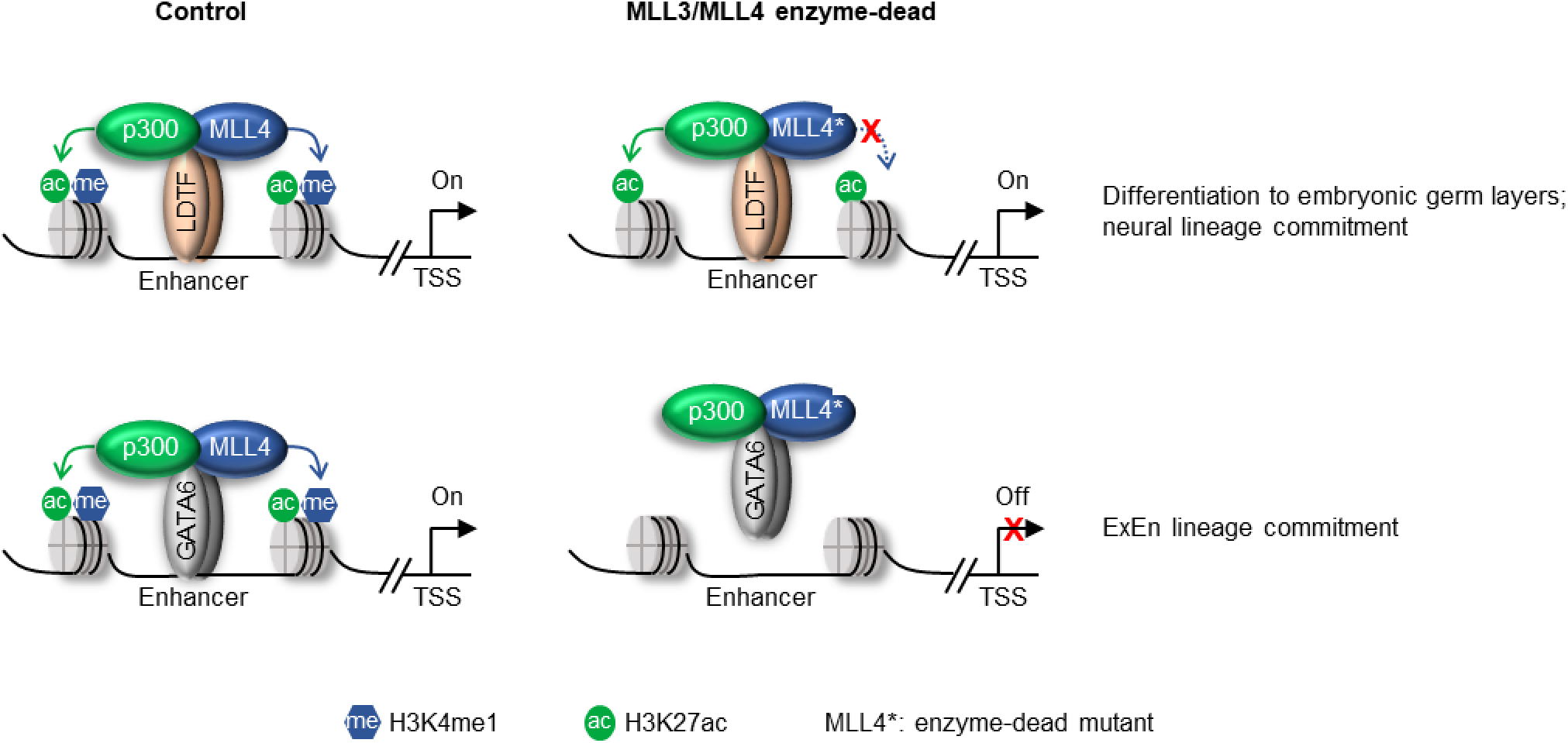
Proposed model. Model depicting lineage-selective but enhancer activation-independent regulation of ESC differentiation and early embryonic development by MLL3/4 methyltransferase activities. LDTF, lineage-determining transcription factor.

The extraembryonic tissues not only provide supportive function, but also play essential roles in patterning the epiblast during early embryonic development^11^. Thus, induction of the PS depends on reciprocal signaling among trophectoderm-derived extraembryonic ectoderm, the ExEn and the epiblast (Fig. 1-S1e)^20, 36^. Without a functional AVE or extraembryonic ectoderm, gastrulation cannot occur. We therefore speculate that gastrulation failure and early embryonic lethality of constitutive dKI embryos are due to defects in extraembryonic tissue development. First, ESCs lacking MLL3/4 enzymatic activities can differentiate towards all three embryonic germ layers. Second, in marked contrast to constitutive dKI embryos which fail in gastrulation, Sox2-dKI embryos develop up to mid-gestation with largely intact gastrulation. Third, constitutive dKI embryos exhibit defective development in VE and fail to form the AVE. Fourth, ESCs lacking MLL3/4 enzymatic activities show severe defects in ExEn differentiation and ectopic GATA6-driven ESC reprogramming to the ExEn lineage. Fifth, MLL3/4 enzymatic activities are required for *Elf5* induction during trophectoderm differentiation. *Elf5* null embryos fail in gastrulation with no signs of PS formation^36^. It is possible that impaired *Elf5* induction and improper trophectoderm development contribute to the failure of PS formation in constitutive dKI embryos. Moreover, defects of cavitation and cardiomyogenesis in KI EBs can be attributed to impaired ExEn induction as reported previously^37, 38^. Supporting this possibility, we found that diminished beating capacity of KI EBs was partially rescued by cultivation with ExEn cell-conditioned medium (Fig. 4-S2f).

Trr is the only homolog of MLL3 and MLL4 in *Drosophila*. A previous study reported that *Drosophila* embryos expressing catalytically deficient Trr by a transgenic rescue method produce fertile adults with only minor abnormalities^39^. However, uniquely in mammals, extraembryonic tissues play essential roles in early embryonic development through tissue interactions and cell signaling^12^. Therefore, mild developmental phenotypes observed in *Trr* mutant flies are not necessarily in conflict with our finding that MLL3/4 enzymatic activities play a major role in extraembryonic tissues rather than embryonic tissues during early mouse development. On the other hand, the mid-gestational lethality of Sox2-dKI embryos suggests that MLL3/4 enzymatic activities are more critical for embryonic development in mammals than in flies.

In undifferentiated ESCs, MLL3/4-catalyzed H3K4me1 has only a minor effect on the maintenance of enhancer activity^9^. In this study, we show for the first time that MLL3/4-catalyzed H3K4me1 is largely dispensable for de novo enhancer activation during cell fate transition using three model systems: EB differentiation, neural differentiation and NEUROD1-driven ESC reprogramming. MLL3/4-catalyzed H3K4me1 has been reported to be required for Cohesin complex binding, enhancer-promoter interaction and gene expression at the *Sox2* locus in ESCs^22^. While our results cast doubt on the general role of H3K4me1-mediated enhancer activation during ESC differentiation, it is possible that H3K4me1-dependent recruitment of Cohesin complex at specific loci precedes and facilitates genomic binding of selective TFs^40^. H3K4 methylation has been reported to antagonize DNA methylation^41^. A recent paper showed that MLL3/4 catalytically-dead ESCs display increased DNA methylation levels on a subset of enhancers^42^. The involvement of DNA methylation in lineage-selective regulation of ESC differentiation by MLL3/4 enzymatic activities remains to be investigated.

We previously reported that MLL3/4 are required for CBP/p300 binding on enhancers activated during cell fate transition^4, 5^. We also provided evidence to suggest that MLL4 protein, rather than MLL4-mediated H3K4me1, controls p300 recruitment to enhancers^4^. In addition, we have shown that MLL4 and BAF complex reciprocally regulate each other’s binding on AEs^43^. Together, these findings offer possible explanations for how MLL3/4 proteins regulate enhancer activation independent of H3K4me1.

## Methods

### Generation of mouse strains

The MLL3 Y4792A and MLL4 Y5477A knockin mouse lines were generated using CRISPR/Cas9 system modified from Wang et al.^44^. Briefly, sgRNA sequences cutting near genomic loci of MLL3 Y4792 (5’-ACTATGGTCATCGAGTACAT-3’) and MLL4 Y5477 (5’-ACGATGGTCATCGAGTACAT-3’) were synthesized by Thermo Fisher’s custom *in vitro* transcription service. The Tyrosine to Alanine point mutations were introduced using single-strand Ultramer DNA Oligonucleotide (IDT). Resulting donor oligonucleotides are as follows: MLL3 Y4792A, 5’-TTCCTTCCACTTAGGGACTGGGCCTGTATGCTGCTAGAGACATTGAAAAACACACTATGGTCATCGA G***GCT***ATTGGAACAATTATTCGAAATGAGGTTGCAAACCGGAAGGAGAAGCTTTATGAGTCTCAGGTA CTGTAT-3’; MLL4Y5477A, 5’-CGCGTATCCAGGGCCTCGGCCTCTATGCAGCCAAGGACCTGGAGAAGCACACGATGGTCATCGAG ***GCT***ATCGGCACCATCATTCGCAATGAGGTGGCCAATCGGCGGGAGAAAATCTATGAGGAGCAGGTA CTGTGGG-3’, in which the inserted mutations are highlighted in italic and bold. To generate each knockin mouse line, sgRNA sequences (20 ng/μL) and its corresponding donor oligonucleotide (100 ng/μL) were co-microinjected with Cas9 nickase mRNA (50 ng/μL, Trilink Biotechnologies) into the cytoplasm of zygotes collected from B6D2F1/J mice (JAX #100006). Injected embryos were cultured in M16 medium (MilliporeSigma) overnight in a 37°C incubator with 6% CO_2_. The next morning, 2-cell-stage embryos were implanted into the oviducts of pseudopregnant surrogate mothers. Offspring were genotyped by PCR and confirmed by Sanger Sequencing.

To generate *Mll3*^KI/KI^;*Mll4*^KI/+^;*Sox2-Cre* mice, *Mll3*^KI/KI^;*Mll4*^KI/+^ or *Mll3*^KI/+^;*Mll4*^KI/+^ mice were crossed with *Sox2-Cre* mice (JAX #008454). *Mll3*^f/f^;*Mll4SET*^f/f^ mice were described previously^10^. *Mll3* ^f/KI^;*Mll4SET* ^f/KI^;*Sox2-Cre* (Sox2-dKI) mice are compound heterozygotes for constitutive *Mll3/4* KI alleles and conditional *Mll3/4* SET domain deletion alleles. The conditional alleles are excised by Cre recombinase in the epiblast and its derivatives, thus resulting in selective elimination of MLL3/4 enzymatic activities in embryonic tissues. All mouse experiments were performed in accordance with the NIH Guide for the Care and Use of Laboratory Animals and approved by the Animal Care and Use Committee of NIDDK, NIH.

### Immunofluorescence and histology

Immunofluorescences were done as previously described^45^. Briefly, mouse blastocysts at E3.5 were flushed out of the uterus in prewarmed M2 medium. Embryos at later developmental stages (E6.5 and E7.5) were collected in cold PBS solution using standard dissection protocols. Blastocysts were fixed with 4% formaldehyde at room temperature for 10 min. Unhatched embryos with intact zona pellucida were digested with Acid Tyrode’s solution prior to fixation. For E6.5 and E7.5, fixation time was extended to 30 min. Next, embryos were washed thrice with PBX (0.1% Triton X-100 in PBS) and permeabilized with 0.5% Triton X-100 and 100mM Glycine in PBS for 5 min for blastocysts and 30 min for E6.5 or E7.5 embryos at room temperature. All incubation and wash steps were performed in polyvinyl clear plates with a thin layer of coating with 1% Agar in 0.9% NaCl to prevent adhesion of embryos. Embryos were incubated with blocking solution (2% horse serum in PBS for blastocysts, 5% horse serum in 0.2% BSA for E6.5 or E7.5) for 2 hours, followed by overnight primary antibody incubation in a humidified chamber covered with a layer of mineral oil to prevent evaporation at 4°C. Next morning, embryos were washed thrice with PBX, incubated again in blocking solution for 2 hours, and transferred into secondary antibody solution for another 2 hours. Blastocysts were incubated at 4°C and E6.5 or E7.5 embryos at room temperature. Following washes in PBX, specimens were counterstained with Hoechst solution (1:5000 in 0.1% PBX), washed again and mounted in Vectashield solution in 8 well chamber slides or 35 mm imaging dishes. Confocal images of embryos were captured using a Zeiss LSM880 laser-scanning microscope. Histology and H&E staining of E18.5 embryos were done as described^3^. In all steps, embryos of the same litters were processed in parallel and imaged with the same conditions and laser intensities.

### Generation of *Mll4* knockin ES cells

The MLL4 Y5477A knockin ES cell lines were generated using the CRISPR/Cas9 system modified from Ran et al.^46^. Briefly, a sgRNA sequence cutting near MLL4 Y5477 (5’-ACGATGGTCATCGAGTACAT-3’) was cloned into lentiCRISPR v2 plasmid (Addgene #52961). The Tyrosine to Alanine point mutation was introduced using single-strand Ultramer DNA Oligonucleotide (IDT). The resulting donor oligonucleotide is as follows: 5’-TCGGCCTCTATGCAGCCAAGGACCTGGAGAAGCACACGATGGTCATCGAG***GCT***ATCGGCACCATCA TTCGCAATGAGGTGGCCAATCGGCGGGAGAAAATCTA-3’, in which the inserted mutations are highlighted in italic and bold. To generate knockin ES cells, the plasmid containing sgRNA (10 μg) and the donor oligonucleotide (400 pmol) were co-transfected into 1 x 10^6^ *Mll3*^-/-^;*Mll4*^f/f^ mouse ES cells by Lipofectamine 3000 (Thermo Fisher). 24 hours later, cells were selected with 1.5 μg/mL puromycin for 3 days. After recovery, 1 x 10^4^ single cells were seeded into a 6 cm dish. Colonies derived from single cells were picked 1 week later and expanded for genotyping. ∼1.1 kb PCR products covering the targeted region were subsequently screened by Sanger sequencing to map the expected mutation. Several potential exonic off-target regions were also checked and no indel was detected. Finally, the stability of the mutated protein was confirmed by immunoblotting.

### ES cell culture

Mouse ES cells were cultured as described in Koehler and Hashino^47^. Briefly, cells were maintained on dishes coated with 0.1% gelatin (Millipore) in 2i+LIF medium, a serum-free ESC culture medium containing 47.75% Advanced DMEM/F-12 (Gibco), 47.75% Neurobasal (Gibco), 0.5% N-2 supplement (Gibco), 1% B-27 supplement minus Vitamin A (Gibco), 1% GlutaMax (Gibco), 1% Penicillin-Streptomycin (Corning), 1% EmbryoMax 2-mercaptoethanol (MilliporeSigma), and supplemented with 1 µM MEK inhibitor PD0325901 (Selleckchem), 3 µM GSK3 inhibitor CHIR99021 HCl (Selleckchem) and 1,000 units/mL mouse LIF protein (MilliporeSigma).

For Alkaline Phosphatase (AP) staining, 4 x 10^3^ single cells were seeded into one well of a 24-well plate. 4 days later, AP staining was done using Alkaline Phosphatase Detection Kit (MilliporeSigma) following the manufacturer’s instructions.

### ES cell differentiation

Embryoid body (EB) differentiation was performed according to the hanging drop method modified from Cao et al.^48^. Briefly, ESCs were suspended into 1 × 10^5^ /mL in indicated differentiation medium and aliquoted into 20 µL drops on the lid of tissue culture dishes. EBs were harvested 2 days later, transferred into ultra-low-attached dishes (Corning) and further cultured in suspension to the indicated time points. The medium used in EB differentiation from day 0 to day 2 is modified from ESC culture medium. It contains no LIF, 1/4 concentration of both inhibitors and is supplemented with 2% ESC-qualified FBS (Gibco). After day 2, the two inhibitors were removed, and medium was changed every other day. The diameter of EB was measured using ImageJ software (NIH). Immunofluorescence of EB was performed as described in Ferguson and Subramanian^49^.

To monitor the cardiomyogenesis during EB differentiation, day 2 EBs harvested from hanging drops were plated into gelatinized 24-well plates with one EB per well. Spontaneous contracting clusters could be observed as early as day 6. The percentage of beating EBs in each 24-well plate was calculated at indicated time points. In the rescue experiment of cardiomyogenesis, KI EBs were cultivated with 50% ExEn cell-conditioned medium or 50% control medium from day 4 to day 6.

Neural differentiation protocol was modified from Bibel et al.^16^. Briefly, neural lineage aggregates were generated following EB differentiation method in ultra-low-attached dishes described above from day 0 to day 8 except that after day 4, 5 µM retinoic acid (MilliporeSigma) was added to the culture. At day 8, aggregates were dissociated with trypsin and neural progenitor cells (NPCs) were plated on PDL/laminin-coated dishes in N-2 medium (Advanced DMEM/F-12 supplemented with 1% N-2 supplement, 1% GlutaMax, 1% EmbryoMax 2-mercaptoethanol and 50 μg/mL BSA). The N-2 medium was changed after 2 hours and again after 1 day. After 2 days, the medium was replaced by the B-27 medium (Neurobasal medium supplemented with 2% B-27 serum free (Gibco), 1% GlutaMax and 1% EmbryoMax 2-mercaptoethanol) to induce neurogenesis. The B-27 medium was changed every other day. Neural differentiation was performed until day 16.

Extraembryonic endoderm (ExEn) differentiation protocol was modified from Ngondo et al.^17^. Briefly, ESCs were seeded at a density of 2 x 10^4^ / cm^2^ into gelatinized plates, and cultured for 5 days in ExEn medium (Advanced RPMI 1640 (Gibco) supplemented with 15% FBS, 1% GlutaMax, 1% EmbryoMax 2-mercaptoethanol, 0.1 μM retinoic acid and 10 ng/mL Activin A (PeproTech)). To collect ExEn cell-conditioned medium, ExEn cells derived from f/f ESCs were washed with PBS twice and resupplied with fresh ExEn medium for 2 days.

Trophectoderm (TE) differentiation protocol was modified from Niwa et al.^28^. Briefly, ESCs were infected with Doxycycline-inducible lentiviral pCW57.1 vector (Addgene #41393) expressing T7-tagged CDX2 or EOMES. Cells were then seeded at a density of 1 x 10^4^ / cm^2^ into gelatinized plates, and cultured for 6 days in TS medium (Advanced RPMI 1640 supplemented with 20% FBS, 1% GlutaMax and 1% EmbryoMax 2-mercaptoethanol) containing 70% MEF-conditioned TS medium and supplemented with 25 ng/mL FGF-4 (MilliporeSigma), 1 μg/mL Heparin (MilliporeSigma) and 1 μg/mL Doxycycline.

To perform GATA6-driven or NEUROD1-driven ESC reprogramming, ESCs were infected with Doxycycline-inducible lentiviral pCW57.1 vector expressing T7-tagged GATA6 or NEUROD1. Cells were then seeded into gelatinize plates and cultured for indicated time periods in serum-containing ESC culture medium (KnockOut™ DMEM (Gibco) supplemented with 15% ESC-qualified FBS, 1% GlutaMax, 1% EmbryoMax 2-mercaptoethanol, 1% HEPES (Gibco), 1% Non-essential AA (MilliporeSigma) and 1,000 units/mL mouse LIF protein) supplemented with 1 μg/mL Doxycycline.

### Teratoma formation assay

Approximately 2 x 10^5^ ESCs were mixed with 50% volume of Matrigel Matrix (Corning) and injected subcutaneously into two sides of an immunocompromised mouse (NSG mouse, JAX #005557). 25 days later, the teratomas were dissected out, fixed in 10% neutral buffered formalin and subjected to histological analysis with H&E staining following standard protocols.

### Immunoblotting and immunoprecipitation

Preparation of nuclear protein extracts and immunoblotting were done as described previously^10^. Briefly, cells were collected and resuspended in cell lysis buffer (10 mM HEPES pH 7.9, 1.5 mM MgCl2, 10 mM KCl and 0.1% NP-40) supplemented with protease inhibitors (Roche) for 20 min at 4 °C to swell cellular membrane. After centrifugation at 2,000x*g*, nuclei were extracted in nuclear lysis buffer (20 mM HEPES, pH 7.9, 1.5 mM MgCl_2_, 420 mM NaCl, 0.2 mM EDTA and 25% glycerol) supplemented with protease inhibitors. Nuclear extracts were resolved using 3–8% Tris-Acetate gradient gels (Invitrogen).

Histone extraction protocol was modified from Shechter et al.^50^. Briefly, cells were collected and resuspended in hypotonic lysis buffer (10 mM Tris-HCl pH 8.0, 1 mM KCl, 1.5 mM MgCl_2_ and 1 mM DTT) supplemented with protease inhibitors for 10 min at 4 °C to swell cellular membrane. After centrifugation at 10,000x*g*, histone fraction was extracted with 0.4 N HCl and neutralized to pH 7.0. Histone extracts were resolved using home-made 12% SDS-PAGE gels.

To extract whole cell lysates, cells or embryos were collected and lysed in RIPA buffer (50 mM Tris-HCl pH 8.0, 150 mM NaCl, 0.1% SDS, 1% NP 40, 0.5% sodium deoxycholate) supplemented with protease inhibitors for 20 min, and subjected to sonication.

Co-immunoprecipitation (IP) was performed as described previously^51^. Briefly, cells were lysed in IP buffer (50 mM Tris-HCl pH 7.5, 150 mM NaCl, 0.3% NP-40 and 2 mM EDTA) supplemented with protease inhibitors for 30 min at 4 °C, and then incubated overnight with Dynabeads Protein A (Thermo Fisher) pre-bound by anti-MLL4 or anti-UTX antibodies. Beads were subsequently washed 5 times using IP buffer. Immunoprecipitated complexes were eluted by boiling the beads for 10 min and resolved using home-made 8% SDS-PAGE gels. All antibodies used in this study are listed in Supplementary Table 2.

### RNA isolation and RT-qPCR analysis

Total RNA was extracted from cultured cells or yolk sac-removed embryos by TRIzol (Life Technologies), and then reverse transcribed using ProtoScript II first-strand cDNA synthesis kit (NEB) following the manufacturer’s instructions. Quantitative PCR was performed in duplicate with Luna Universal qPCR Master Mix (NEB) using QuantStudio 5 Real-Time PCR System (Thermo Fisher). RT-qPCR data were normalized using *Gapdh* and were presented as means ± SD. Primer sequences used in RT-qPCR are listed in Supplementary Table 3.

### RNA-seq library preparation

1 µg total RNA was used for mRNA purification with NEBNext Poly(A) mRNA Magnetic Isolation Module (NEB, E7490). From purified mRNA, libraries were constructed using NEBNext Ultra™ II RNA Library Prep Kit for Illumina (NEB, E7770) following the manufacturer’s instructions, and were sequenced on Illumina HiSeq 4000 and NovaSeq 6000.

### ChIP and ChIP-seq library preparation

Chromatin immunoprecipitation (ChIP) and ChIP-Seq were performed as described previously^4^. Briefly, cells were cross-linked with 1% formaldehyde for 10 min and quenched by 125 mM glycine for 10 min. Fixed cells were swelled in the lysis buffer containing 5mM PIPES pH 7.5, 85mM KCl, 0.5% NP-40 and protease inhibitors, incubated on ice for 20 min, and centrifuged at 500x*g* for 5 min at 4°C. Nuclei were resuspended with cold TE buffer (10mM Tris-HCl pH8.0, 1mM EDTA), and subjected to sonication. Sheared chromatin was clarified by centrifugation at 13,000x*g* for 10 min at 4°C. The supernatant was transferred to a new tube and further supplemented with 150mM NaCl, 1% Triton X-100, 0.1% sodium deoxycholate and protease inhibitors. 2% of the mixture was set aside as input. For each ChIP, chromatin from 1 x 10^7^ cells were mixed with 20 ng spike-in chromatin (Active Motif, #53083) and incubated with 4-8 μg primary target antibody and 2 μg spike-in antibody (Active Motif, #61686) overnight at 4°C. Then, ChIP samples were precipitated by 50 μL prewashed Dynabeads Protein A. Beads were washed twice with 1 mL RIPA buffer, once with 1 mL RIPA buffer containing 500mM NaCl, twice with 1 mL LiCl buffer and once with 1 mL TE buffer, and then eluted with 100 μL fresh elution buffer (0.1M NaHCO_3_, 1% SDS). ChIP samples and input were incubated with Proteinase K (NEB) at 65°C overnight to reverse formaldehyde crosslinking, and then purified using QIAquick PCR purification kit (Qiagen).

For ChIP-Seq, the entire ChIP DNA and 300 ng input DNA were used to construct libraries using NEBNext Ultra™ II DNA Library Prep kit for Illumina (NEB, E7645) following the manufacturer’s instructions. The final libraries were sequenced on Illumina HiSeq 3000 and HiSeq 4000.

### ATAC-seq library preparation

Assay for Transposase-Accessible Chromatin with high-throughput sequencing (ATAC-Seq) was performed as described^52^. Briefly, for each reaction, 5 × 10^4^ cells were freshly collected, washed and swelled in 50 µL cold lysis buffer (10 mM Tris-HCl pH 7.5, 10 mM NaCl, 3 mM MgCl_2_, 0.1% NP-40, 0.1% Tween-20, and 0.01% digitonin) for 3 min. Nuclei were collected by centrifugation at 500×*g* for 10 min at 4°C, and then resuspended in 50 µL of transposition reaction buffer containing 25 µL 2x Tagment DNA buffer, 2.5 µL transposase (100 nM final, Illumina), 0.5 µL 1% digitonin, 0.5 µL 10% Tween-20, 16.5 µL PBS, and 5 µL H_2_O. The reaction was mixed for 30 min at 37°C and subjected to purification using the MinElute Reaction Cleanup Kit (Qiagen) according to the manufacturer’s instructions. Purified DNA was amplified for library construction with PCR using Nextera i5 common adaptor and i7 index adaptors (Illumina). The final libraries were sequenced on Illumina HiSeq 4000.

### Computational analysis

#### NGS data processing

For RNA-seq, raw sequencing data were aligned to the mouse genome mm9 using STAR software^53^ (v2.7.5). Reads on exons defined by UCSC gene annotation system were collected to calculate Transcripts Per Million (TPM) as a measure of gene expression level, and genes with TPM > 3 were regarded as expressed.

For ChIP-seq, raw sequencing data were aligned to the mouse genome mm9 and the drosophila genome dm6 using Bowtie2^54^ (v2.3.2). To identify ChIP-enriched regions, SICER^55^ (v1.1) was used. For ChIP-Seq of GATA6, MLL4 and T7 tag, the window size of 50 bp, the gap size of 50 bp, and the false discovery rate (FDR) threshold of 10^-10^ were used. For ChIP-Seq of histone modifications (H3K4me1 and H3K27ac), the window size of 200 bp, the gap size of 200 bp, and the FDR threshold of 10^-3^ were used. Reads on indicated regions were collected to calculate Reads Per Kilobase Million (RPKM) as a measure of signal intensity.

For ATAC-seq, raw ATAC-Seq reads were processed using Kudaje lab’s ataqc pipeline (https://github.com/kundajelab/atac_dnase_pipelines) (v0.3.4) which includes adapter trimming, aligning to mouse mm9 genome by Bowtie2, and peak calling by MACS2^56^. For downstream analysis, filtered reads that were retained after removing unmapped reads, duplicates and mitochondrial reads were used.

#### RNA-seq analysis

Differentially expressed genes were determined using DESeq2^57^ (v1.20.0) in R/Bioconductor with a cutoff of 2.5-fold (adjusted *p* < 0.05). PCA graph, volcano plot and scatter plots were drawn using ggplot2^58^ (v3.2.1) in R. Heat maps were drawn using pheatmap^59^ (v1.0.12) in R. Gene ontology (GO) analysis was done using DAVID^60^ with the whole mouse genome as background.

Gene set enrichment analysis (GSEA) was performed using GSEA software^61^ (v4.0.3). Cell-type markers in E6.5 embryos with *p* < 0.05^24^ were used as signature databases in Fig. 4-S1a.

#### Analysis of ChIP-seq peaks

To define regulatory regions, a combination of genomic coordinates and histone modification ChIP-Seq data were used. Promoter regions were defined as transcription start sites ± 1 kb. Promoter-distal regions were further overlapped with H3K27ac^+^ regions to define active enhancers (AEs). Target^+^ AEs were defined as AEs that are overlapping with target peaks for at least 1 bp.

In Fig. 4e, de novo AEs were defined as AEs with over 2-fold H3K27ac intensity increase in f/f cells after ExEn differentiation compared to in ESCs. In Fig. 5d,h, GATA6-T7 or NEUROD1-T7 induced AEs were defined as GATA6-T7^+^ or NEUROD1-T7^+^ AEs with over 2-fold H3K27ac intensity increase in cells after 1-day Doxycycline treatment compared to in undifferentiated ESCs. In Fig. 7-S1b, all MLL4^+^ AEs existed in D4 EBs but not in ESCs were regarded as de novo MLL4^+^ AEs. To compare MLL4 genomic distributions between f/f and KI D4 EBs, the union of de novo MLL4^+^ AEs was divided into three groups based on MLL4 binding intensities in KI cells compared to f/f cells with a cutoff of 2-fold.

Heat map matrices were generated using in-house scripts with 50 bp resolution and visualized in R. Average profiles were plotted using the number of ChIP-seq reads (normalized to the size of each library) in 5 bp bins from the center of each regulatory element to 2 kb on both sides.

In Fig. 4g, induced genes associated with GATA6^+^ MLL4^+^ de novo AEs were identified by assigning each GATA6^+^ MLL4^+^ de novo AE to its closest genes induced during ExEn differentiation (TSSs in ±100 kb). In Fig. 7-S1e, we associated de novo MLL4^+^ AEs from each group to all genes expressed in D4 EBs (TSSs in ±100 kb), and calculated a regulatory potential score for each associated gene as the sum of the contribution of individual AEs from this group using BETA^62^. The gene associated predominantly with a certain group of de novo MLL4^+^ AEs was identified as its regulatory potential score in this group is higher than that in any of the other 2 groups.

#### Motif analysis

To find enriched TF motifs in given genomic regions identified by ChIP-Seq, we utilized the SeqPos motif tool in the Cistrome toolbox^63^. For ChIP-Seq data with more than 5,000 regions, top 5,000 significant regions were used. For ChIP-Seq data with small numbers, all regions were used.

#### Analysis of super-enhancer

We used Rank Ordering of Super-Enhancers^64^ (ROSE) with default parameters to identify super-enhancers (SEs). To identify SEs in NPCs, we stitched SOX3 binding sites (GSE33059)^35^ in NPC AEs and used H3K27ac signal intensity for ranking. Genes associated with NPC SEs were identified by assigning each NPC SE to its closest genes (TSSs in ±100 kb).

#### Box plots and genomic profile visualization

Box plots were drawn with GraphPad Prism 9 software using the Tukey method. In Fig. 7e-f, ChIP-seq signal intensities of MLL4, H3K4me1 and H3K27ac on NPC SEs, and RNA-seq defining expression levels of NPC SE-associated genes in f/f and KI cells were plotted. In Fig. 7-S1e, RNA-seq defining expression levels of genes associated predominantly with each of the 3 groups of de novo MLL4^+^ AEs were calculated as fold changes (log_2_) and were plotted. Wilcoxon signed-rank test (two-sided) was used to determine statistical differences in signal enrichment and gene expression levels between f/f and KI cells.

To visualize data from ChIP-seq, ATAC-seq and RNA-seq in the Integrative Genomics Viewer^65^ (IGV) (v2.4.4), reads were collected and converted to wiggle (wig) format-profiles using in-house script.

## Acknowledgement

We thank Younghoon Jang for assistance in generating *Mll3* KI mice, Kaitlin McKernan and Ruth Kopyto for assistance in genotyping, Danyang Wan and Hui Sun for assistance in EB immunofluorescence, Anna-Katerina Hadjantonakis and Andrew Xiao for suggestions on mouse experiments, Hualong Yan for suggestions on ESC culture, and all members of the Ge laboratory for discussions and support. We also thank NHLBI DNA Sequencing and Genomics Core and UCSD IGM Genomics Center for next-generation sequencing, NIH HPC group for high-performance computing systems. This work was supported by the Intramural Research Program of NIDDK, NIH to K.G.

## Author Contributions

G.X. and K.G. conceived the project. G.X., J.-E.L., W.P. and C.L. performed the methodology. G.X., A.D.S., Y.-K.P., S.C. and J.J.T. carried out the experiments. J.-E.L., G.X. and W.P. performed bioinformatics analyses. G.X. and K.G. wrote the manuscript with inputs from all authors. K.G., P.P.R and T.S.M provided supervision and acquired funding.

## Declaration of Interests

The authors declare no competing interests.

## Data availability

All data sets described in the paper have been deposited in NCBI Gene Expression Omnibus under accession number GSE154475.

## Code availability

In-house generated computational codes are available upon request.

## Supplementary Information

Supplementary Table 1: Detailed information of mating experiments

Supplementary Table 2: Information of antibodies

Supplementary Table 3: Sequences of RT-qPCR primers

---

**Fig. 1-S1:**
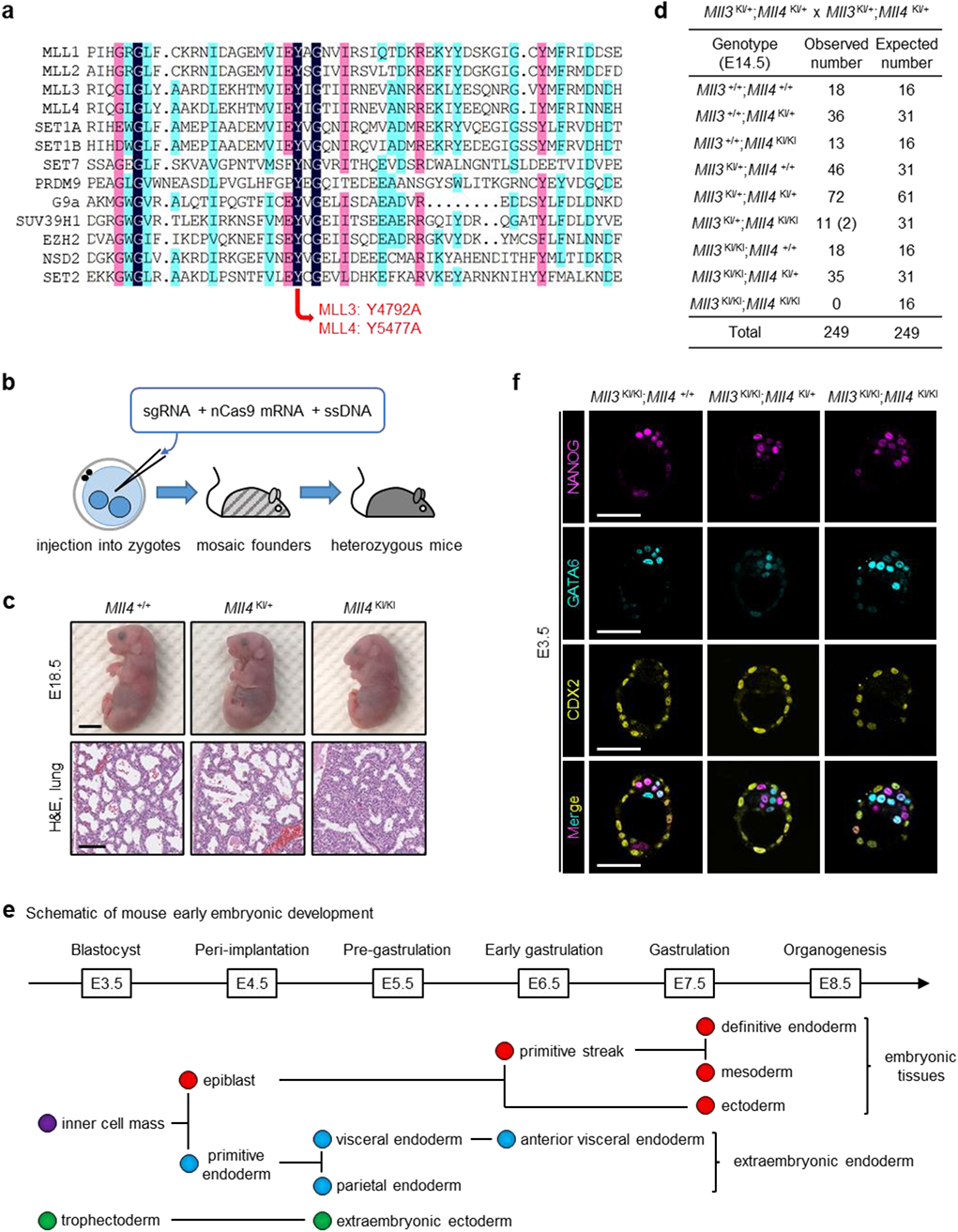
Loss of MLL3/4 enzymatic activities results in embryonic lethality. **a**, Alignment of SET domain sequences in different histone lysine methyltransferases in mice. The highly conserved tyrosine (Y) residues, Y4792 in MLL3 and Y5477 in MLL4, are mutated to alanine residues (A) in mice and/or ESCs in this study. **b**, Schematic of generating enzyme-dead MLL3/MLL4 knockin (KI) mice by Cas9 nickase. sgRNA, single guide RNA; nCas9, Cas9 nickase; ssDNA, single-strand DNA. **c**, Representative images of embryos with indicated genotypes and H&E-stained lung tissues at E18.5. Scale bar, 5 mm (embryo) or 250 μm (histology). **d**, Genotypes of progeny from *Mll3*^KI/+^;*Mll4*^KI/+^ intercrosses at E14.5. Numbers of dead pups are indicated in parentheses. **e**, Representative images of NANOG, GATA6 and CDX2 IF staining of E3.5 blastocysts. Scale bar, 100 μm. **f**, Schematic of early embryonic development in mice.

**Fig. 2-S1:**
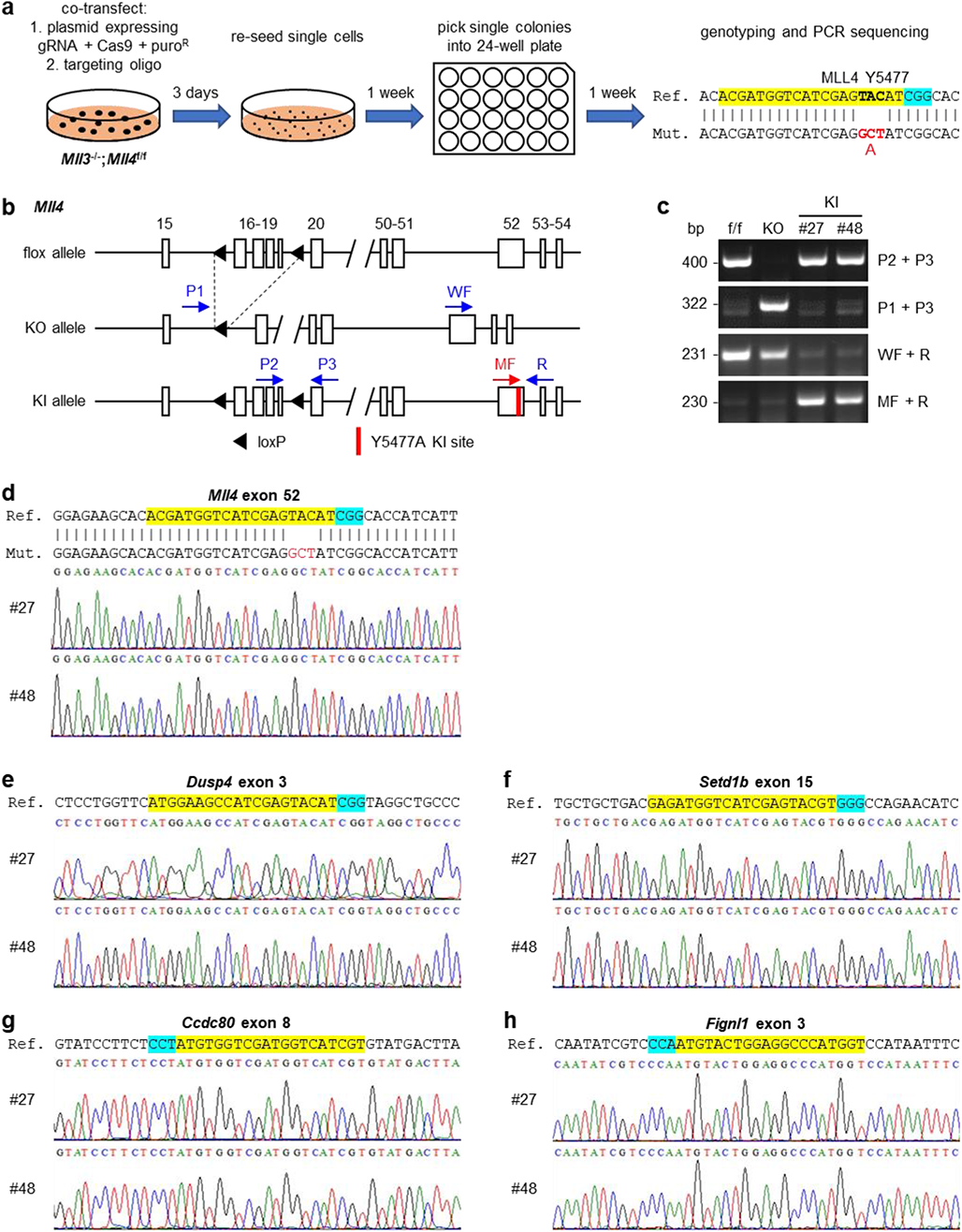
Generation of ESCs lacking MLL3/4 enzymatic activities. **a**, Schematic of generating ESCs harboring enzyme-dead mutant (Y5477A) MLL4. Knockin was done in *Mll3*^-/-^;*Mll4*^f/f^ ESCs. **b**, Schematic representation of *Mll4* flox, knockout (KO) and knockin (KI) alleles. In flox and KI alleles, two loxP sites were inserted in the intron before exon 16 and the intron after exon 19; in the KI allele, the Y5477A mutation is located in exon 52. Locations of PCR genotyping primers P1, P2, P3, WF, MF and R are indicated by arrows. The MF primer is specific to the KI allele. **c**, PCR genotyping using primer pairs indicated in **b**. Sizes of PCR products are indicated on the left. **d-h**, Chromatograms of genomic PCR sequencing from two independent KI ES cell lines. Trace files display sequences around the Y5477A mutation (**d**) or sequences around potential exonic off-target regions of *Dusp4* (**e**), *Setd1b* (**f**), *Ccdc80* (**g**) and *Fignl1* (**h**). In the reference sequences, sgRNA-defined target regions and PAM sites are highlighted in yellow and cyan, respectively.

**Fig. 3-S1:**
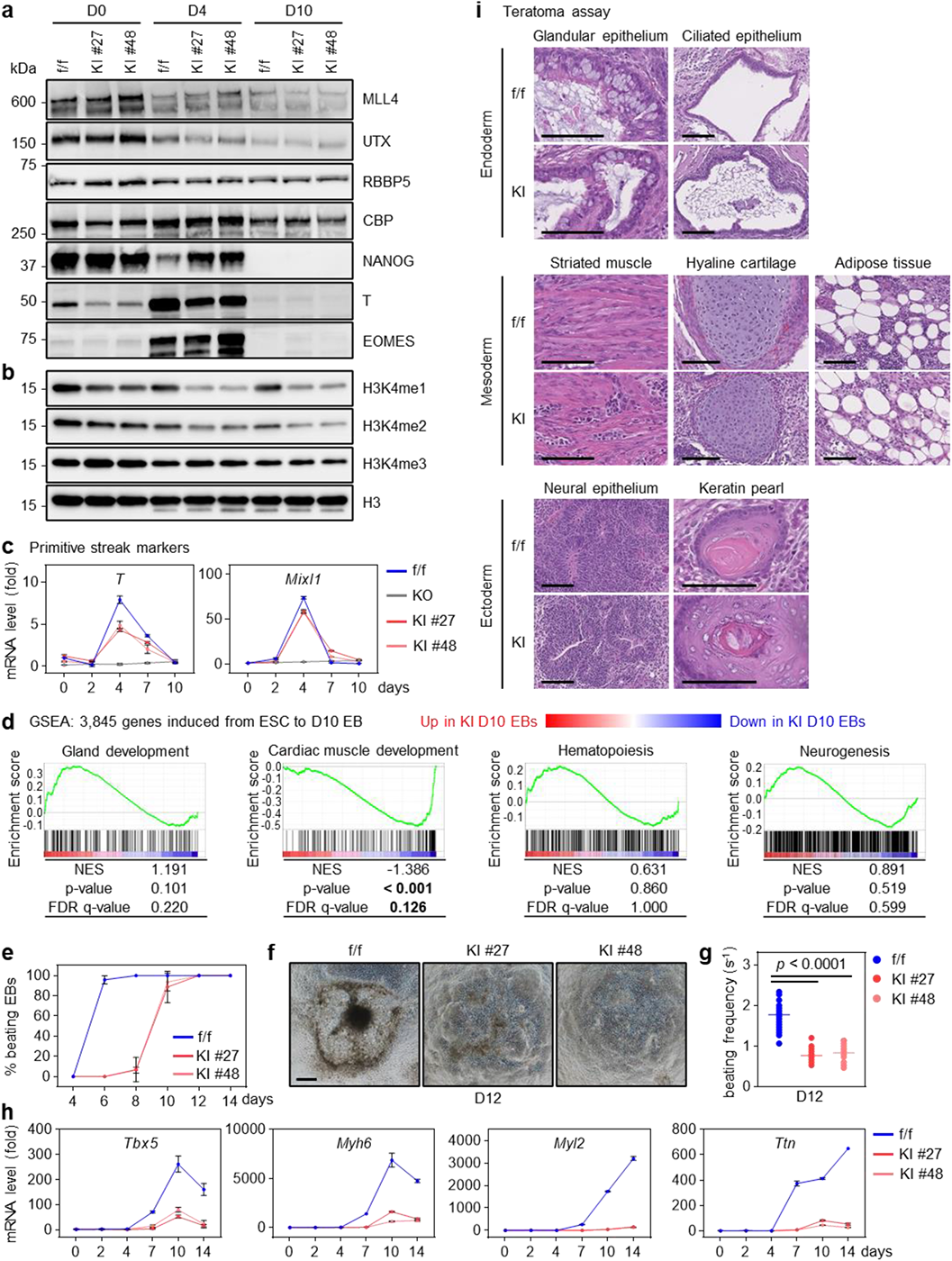
ESCs lacking MLL3/4 enzymatic activities can differentiate towards the three germ layers, but show defects in cardiomyogenesis. **a**,**b**, Immunoblotting in f/f and KI cells at day 0 (D0), D4 and D10 of EB differentiation. Nuclear extracts (**a**) or histone extracts (**b**) were analyzed using indicated antibodies. **c**, RT-qPCR analysis of primitive streak markers (*T*, *Mixl1*) at indicated time points. **d**, Gene set enrichment analysis (GSEA) of 3,845 genes induced from ESC to D10 EB on developmental terms. NES, normalized enrichment score. Statistically significant data are highlighted in bold. **e**, The percentage of beating EBs at indicated time points. The result is from 3 independent experiments with 24 EBs per group. **f**, Representative microscopic images of attached EBs at D12. Scale bar, 250 μm. **g**, Beating frequencies of EBs at D12 are presented as dot plots. Horizontal lines represent mean values. 24 EBs were measured per group. Statistical significance was determined by the two-tailed unpaired *t*-test. **h**, RT-qPCR analysis of the cardiac progenitor marker (*Tbx5*) and cardiac myofilament genes (*Myh6*, *Myl2*, *Ttn*) at indicated time points. **i**, Teratoma assay of f/f and KI ESCs. Representative histological sections of structures belonging to endoderm, mesoderm and ectoderm are shown. Scale bar, 100 μm.

**Fig. 3-S2:**
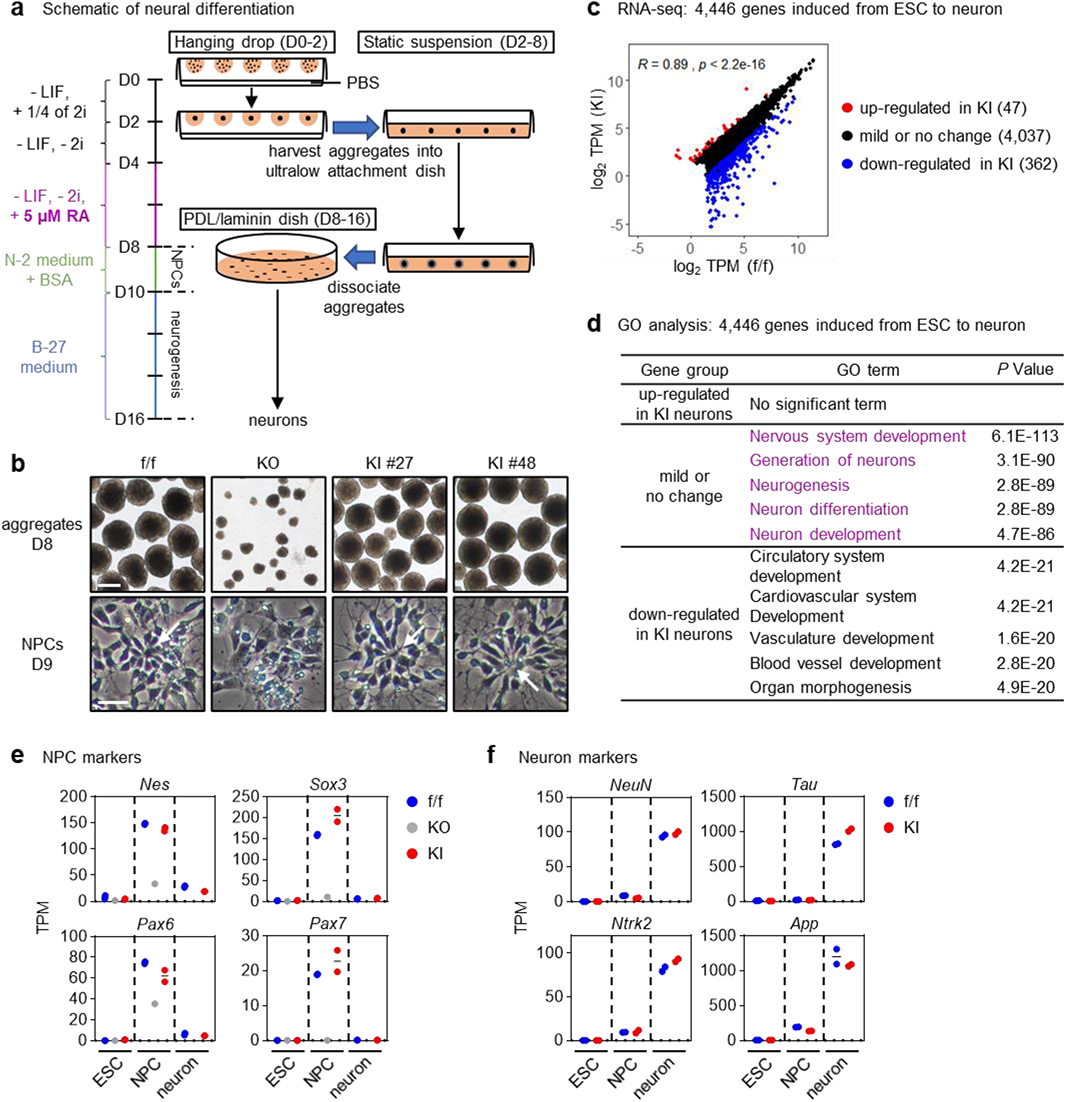
MLL3/4 enzymatic activities are generally dispensable for neural differentiation. **a**, Schematic of neural differentiation. RA, Retinoic acid. **b**, Representative microscopic images of D8 aggregates and D9 NPCs. Scale bar, 250 μm (*upper*) or 25 μm (*lower*). White arrows indicate neural rosettes. **c**, Expression levels of 4,446 genes induced from ESC to neuron were presented as scatter plots. Pearson correlation coefficient (*R*) and *P* value are shown. The number of genes in each group is indicated in parentheses. **d**, GO analysis of three gene groups in **e**. Terms associated with neural differentiation are highlighted in purple. **e**,**f**, Expression levels of NPC (**e**) and neuron (**f**) markers. Data from RNA-seq are presented as dot plots (f/f, n = 2; KI, n = 2). Horizontal lines represent mean values. **e**, Data of KO ESCs and NPCs from RNA-seq are included. **g**,**h**, Violin plots of expression levels of genes induced from ESC to NPC (**g**) or from ESC to neuron (**h**). Induced genes were grouped based on differential expression between f/f and KI cells. Center lines represent median values; dash lines represent lower and upper quartiles. Statistical significance was determined by the one-sided Mann-Whitney test.

**Fig. 4-S1:**
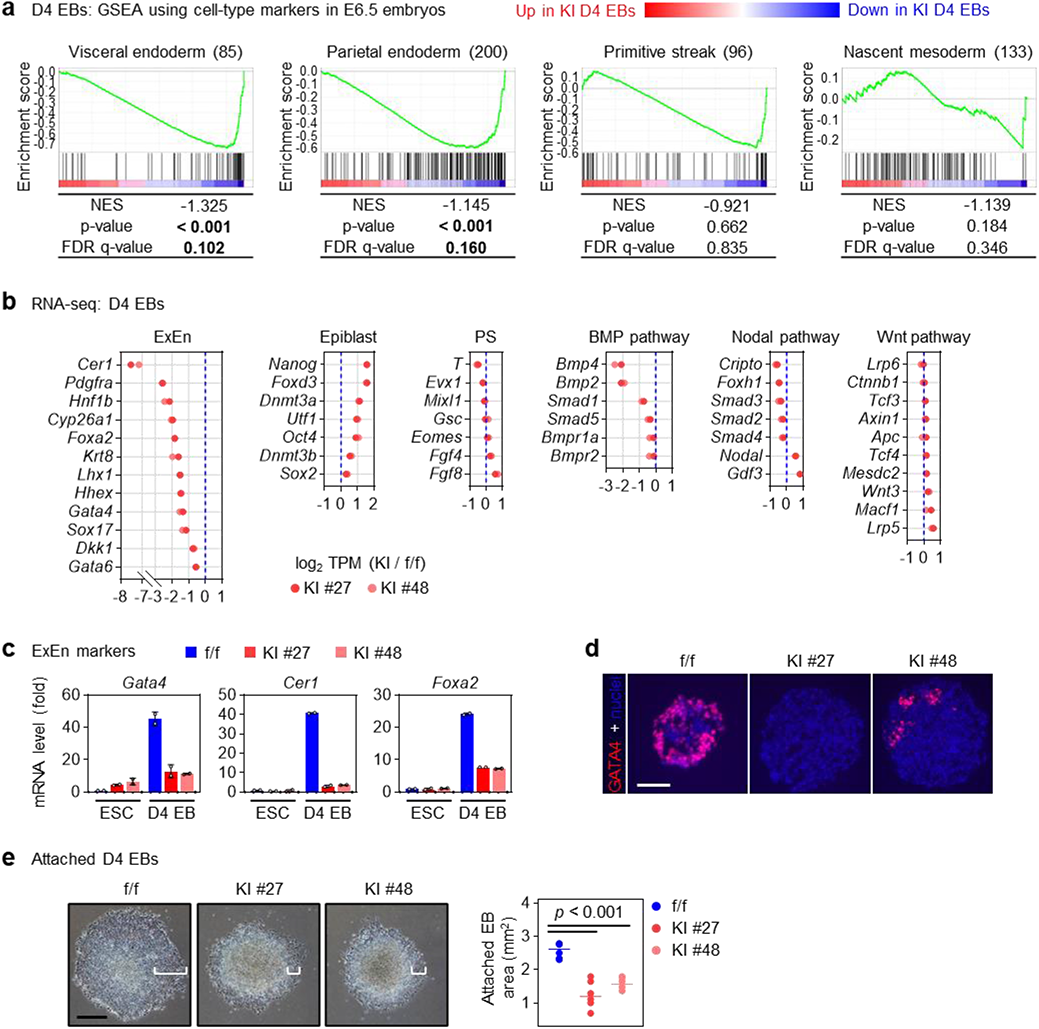
Loss of MLL3/4 enzymatic activities impairs ExEn gene induction during EB differentiation. **a**, GSEA of expression profiles in D4 EBs using cell-type markers defined in E6.5 embryos^24^. NES, normalized enrichment score. Statistically significant data are highlighted in bold. **b**, Expression fold changes of markers of extraembryonic endoderm (ExEn), epiblast and primitive streak (PS), as well as of genes encoding components of BMP, Wnt and Nodal signaling pathways, between KI and f/f D4 EBs. **c**, ExEn markers were analyzed by RT-qPCR in ESCs and D4 EBs (*n* = 2). **d**, IF staining of GATA4 in D4 EBs. Scale bar, 100 μm. **e**, D2 EBs in suspension were plated onto gelatinized surfaces. Representative microscopic images of attached EBs at D4 are shown (*left*). ExEn-like cells migrating away from the EB periphery are indicated with brackets. Scale bar, 250 μm. Areas of attached EBs at D4 are presented as dot plots (*right*). Horizontal lines represent mean values. 6 EBs were measured per group. Statistical significance was determined by the two-tailed unpaired *t*-test.

**Fig. 4-S2:**
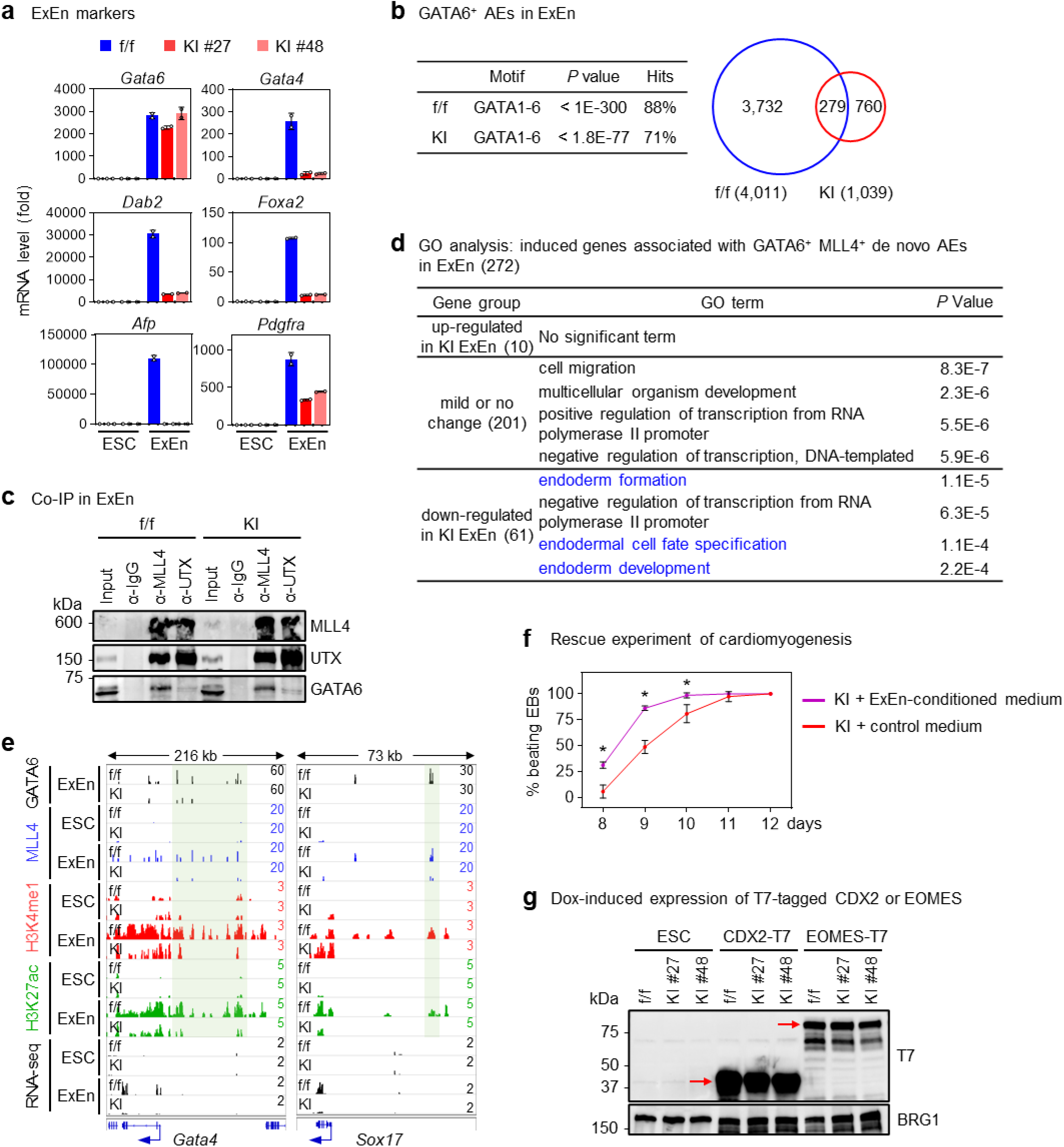
ESCs lacking MLL3/4 enzymatic activities show aberrant differentiation to extraembryonic lineages. **a**, ExEn markers were analyzed by RT-qPCR before and after ExEn differentiation (*n* = 2). **b**, Motif analysis and Venn diagram of GATA6^+^ AEs in f/f and KI cells after ExEn differentiation. **c**, Whole cell lysates prepared from cells after ExEn differentiation were immunoprecipitated with MLL4 or UTX antibodies. Immunoprecipitated complexes were analyzed by immunoblotting using antibodies against MLL4, UTX and GATA6. **d**, GO analysis of three gene groups in Fig. 4g. Terms associated with ExEn differentiation are highlighted in blue. **e**, ChIP-seq profiles of GATA6, MLL4, H3K4me1 and H3K27ac as well as RNA-seq profiles in f/f and KI cells are displayed on *Gata4* and *Sox17* loci. GATA6^+^ MLL4^+^ de novo AEs are highlighted in shades. **f**, During EB differentiation, KI cells were cultivated with ExEn cell-conditioned medium or control medium. The percentage of beating EBs at indicated time points were shown. The result is from 3 independent experiments with 24 EBs per group. **g**, Immunoblotting of T7-tagged CDX2 and EOMES. f/f and KI ESCs were infected with Doxycycline (Dox)-inducible lentiviral vector expressing T7-tagged CDX2 or EOMES and treated with 1 μg/ml Dox for 1 day. Whole cell lysates were analyzed using indicated antibodies. BRG1 is shown as loading control.

**Fig. 6-S1:**
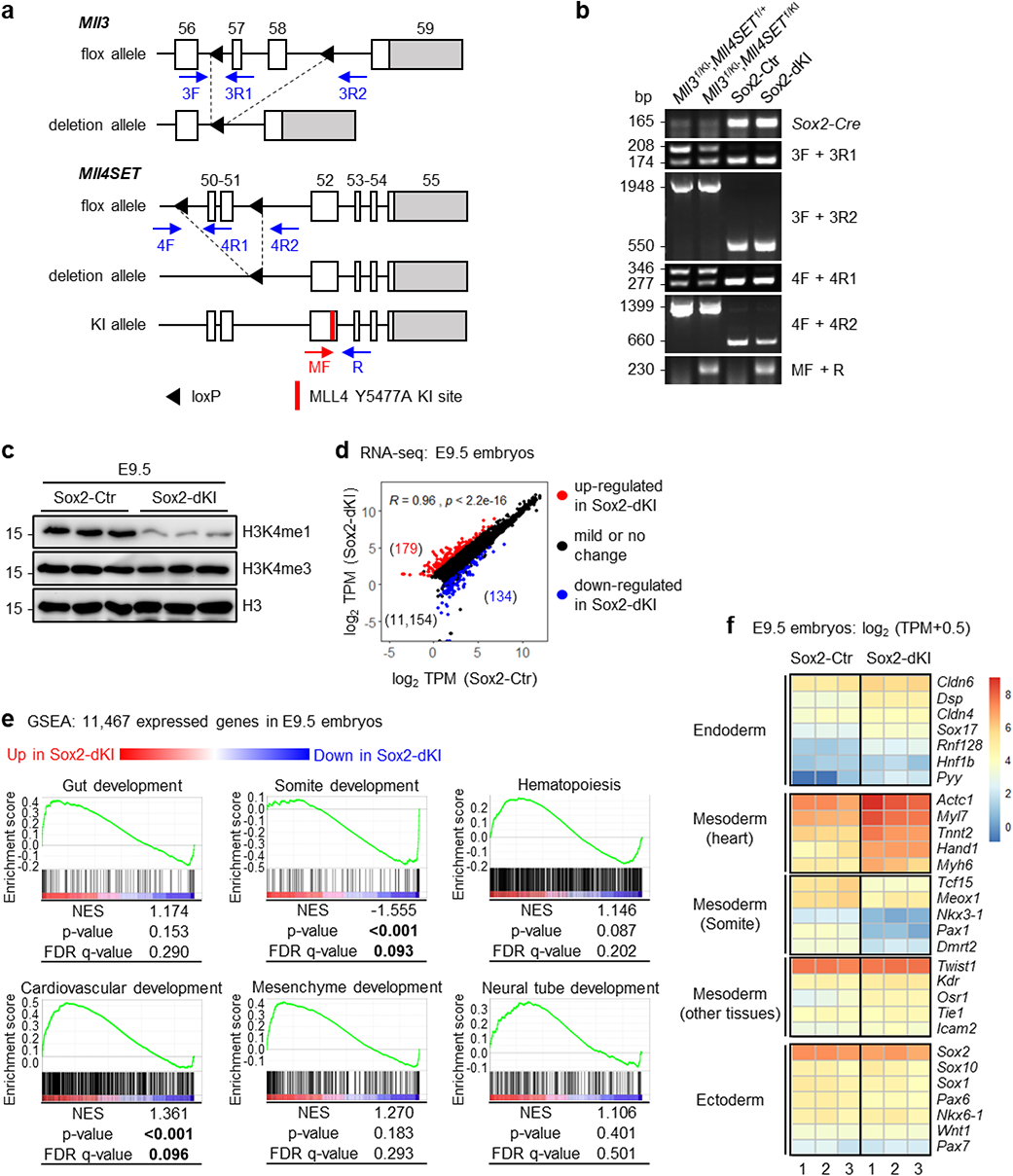
Lineage-selective roles of MLL3/4 enzymatic activities in early embryonic development. **a**, Schematic representation of *Mll3* flox and deletion, as well as *Mll4SET* flox, deletion and KI alleles^10^. Locations of PCR genotyping primers are indicated by arrows. The MF primer is specific to the KI allele. **b**, PCR genotyping of E9.5 yolk sac-removed embryos using primers showed in Fig. 6-S1a as well as primers detecting *Sox2-Cre* allele. Sizes of PCR products are indicated on the left. **c**, Whole cell lysates prepared from E9.5 yolk sac-removed embryos were analyzed with immunoblotting using indicated antibodies. **d**, Sox2-Ctr and Sox2-dKI embryos were collected at E9.5 for RNA-seq. Levels of expressed genes were presented as scatter plots. Pearson correlation coefficient (*R*) and *P* value are shown. The number of genes in each group is indicated in parentheses. **e**, GSEA of 11,467 expressed genes in E9.5 embryos on developmental terms. NES, normalized enrichment score. Statistically significant data are highlighted in bold. **f**, Heat maps of gene expression level in E9.5 embryos for markers of germ layer derivatives. Data from 3 embryos were presented for each genotype.

**Fig. 7-S1:**
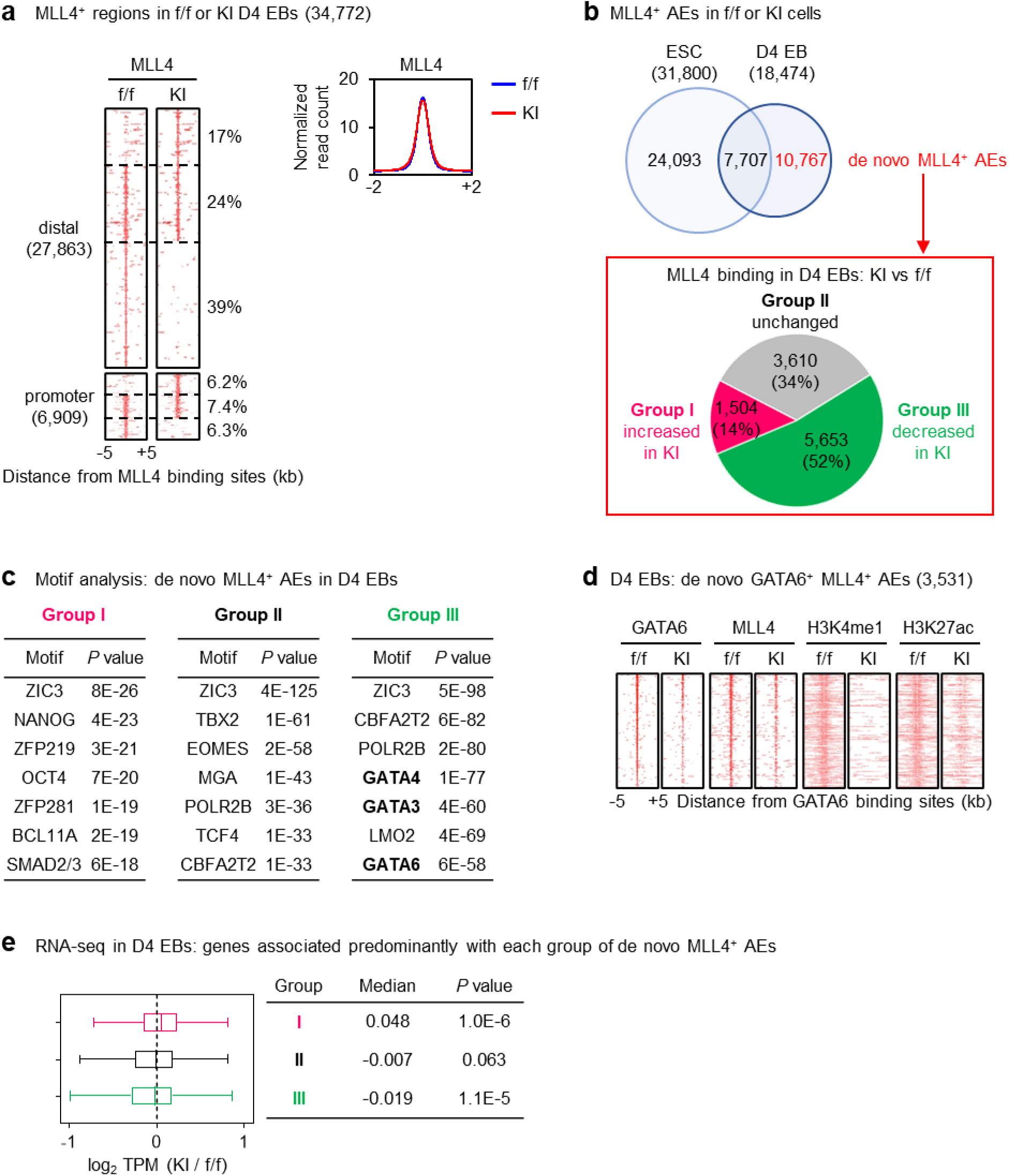
Loss of MLL3/4 enzymatic activities results in redistribution of MLL4 genomic binding. **a**, D4 EBs were collected for ChIP-seq of MLL4. 34,772 MLL4^+^ regions were identified in f/f or KI ESCs. MLL4^+^ distal regions or promoters were split based on the MLL4 binding intensities in KI EBs compared to f/f EBs. Average profiles (*left*) and heat maps (*right*) of 34,772 MLL4^+^ regions in f/f and KI EBs are shown. **b**, MLL4^+^ AEs in ESCs and D4 EBs are depicted by the Venn diagram. 10,767 de novo MLL4^+^ AEs were divided into three groups based on the changes of MLL4 binding intensities from f/f to KI EBs: increased (Group I), unchanged (Group II) and decreased (Group III). **c**, Motif analysis of the three groups of de novo MLL4^+^ AEs. GATA family transcription factors were highlighted in bold. **d**, Heat maps of GATA6 and MLL4 genomic bindings as well as H3K4me1 and H3K27ac enrichments on de novo GATA6^+^ MLL4^+^ AEs in f/f and KI D4 EBs. **e**, Expression fold changes (log_2_) of genes associated predominantly with each group of de novo MLL4^+^AEs. Data from RNA-seq are presented in box plots. Center lines represent median values; the bottom and top of the boxes represent lower and upper quartiles; whiskers were calculated using the Tukey method. Statistical significance was determined by the two-sided Wilcoxon signed-rank test.

**Fig. 7-S2:**
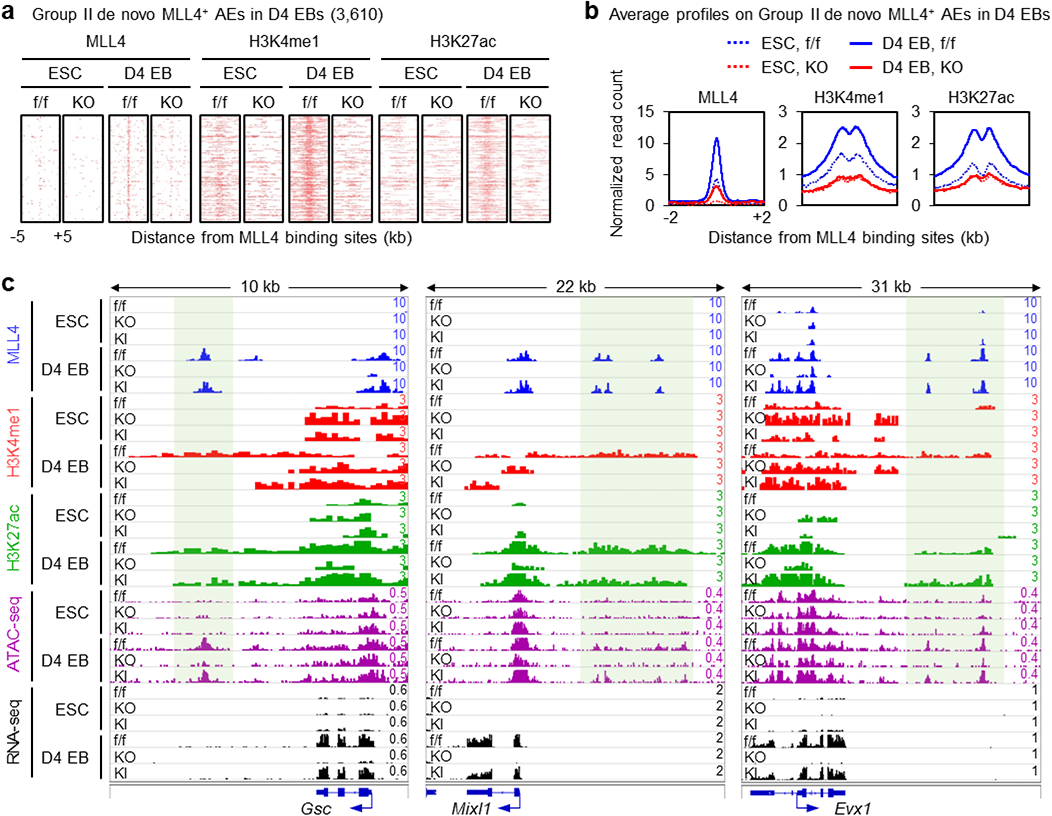
MLL3/4 proteins, but not MLL3/4-catalyzed H3K4me1, are required for activation of Group II AEs during early EB differentiation. **a**,**b**, Heat maps (**a**) and average profiles (**b**) of MLL4 genomic bindings, H3K4me1 and H3K27ac enrichments in f/f and KO cells on Group II de novo MLL4^+^ AEs identified in Fig. 7-S1b. ChIP-seq data in KO cells were obtained from GSE50534^4^ and reanalyzed. **c**, ChIP-seq profiles of the same datasets as in Fig. 7c are displayed on *Gsc*, *Mixl1* and *Evx1* loci. Group II de novo MLL4^+^ AEs are highlighted in shades.

**Fig. 7-S3:**
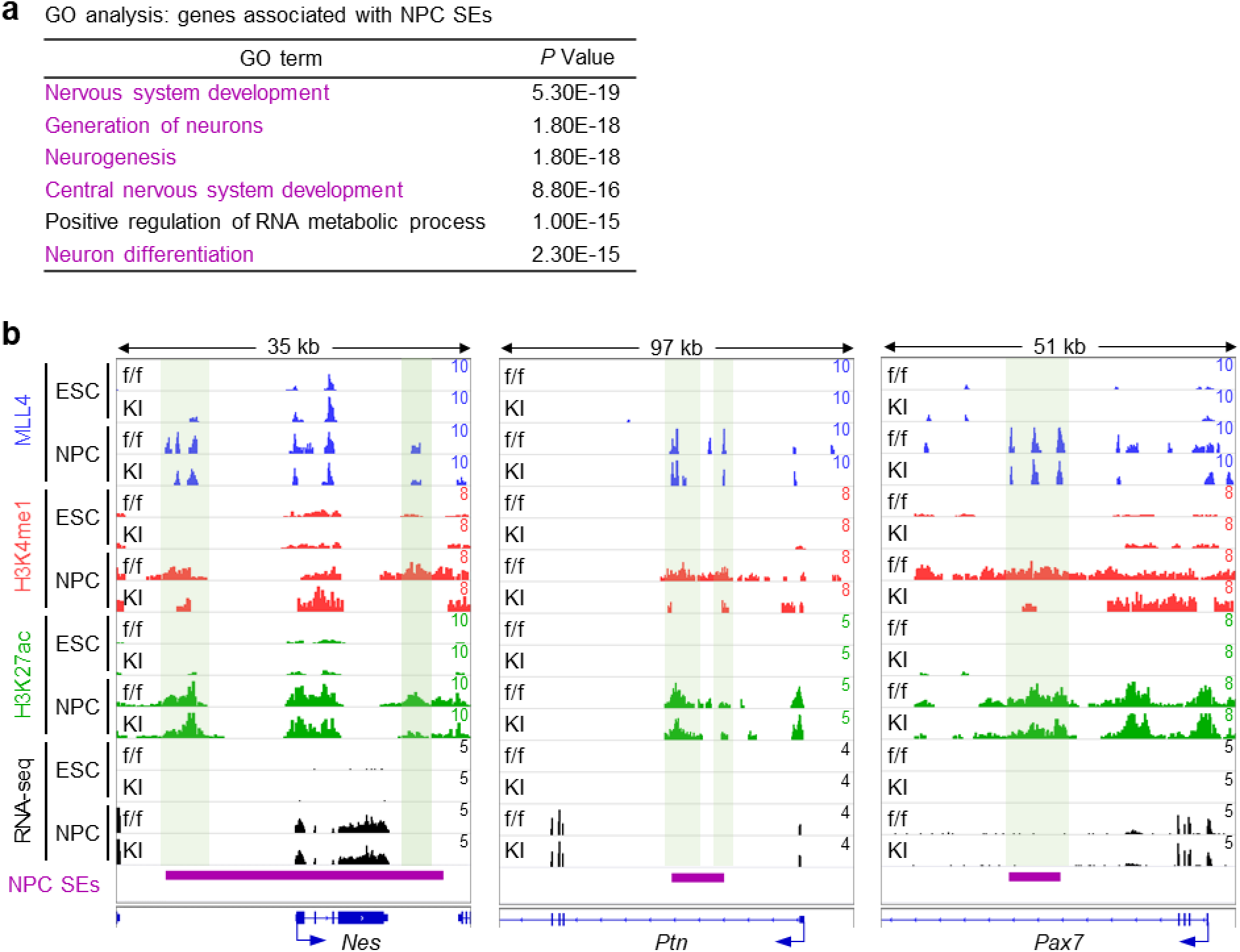
MLL3/4-catalyzed H3K4me1 is dispensable for activation of super-enhancers during neural differentiation. **a**, GO analysis of genes associated NPC SEs. Terms associated with neural differentiation are highlighted in purple. **b**, ChIP-seq profiles of MLL4, H3K4me1 and H3K27ac, and RNA-seq profiles in f/f and KI cells are displayed on *Nes*, *Ptn* and *Pax7* loci. NPC SEs are indicated by purple bars and de novo MLL4^+^ enhancers in SEs are highlighted in shades.

## Notes

### Competing Interest Statement

The authors have declared no competing interest.

### Summary of Updates

1. New Fig. 1e-g to show that MLL3/4 enzymatic activities are essential to initiate gastrulation during early embryonic development of mice. 2. New Fig. 4 to show that loss of MLL3/4 enzymatic activities leads to aberrant ESC differentiation to extraembryonic lineages including extraembryonic endoderm (ExEn) and trophectoderm. 3. New Fig. 5 to show that MLL3/4 enzymatic activities are required for GATA6 binding on enhancers, which provides a plausible explanation for the ExEn differentiation defects of KI ESCs. 4. New Fig. 6 to show that consistent with ESC data, Sox2-Cre-mediated selective elimination of MLL3/4 enzymatic activities in embryonic tissues leaves gastrulation largely intact. 5. In summary, we propose a model in the new Fig. 8 that MLL3/4 enzymatic activities regulate ESC differentiation and early embryonic development through selectively modulating genomic binding of lineage-determining transcription factors, rather than directly activating enhancers. 6. Based on our new findings, we have changed the title of this manuscript.

